# Activation of polarized cell growth by inhibition of cell polarity

**DOI:** 10.1101/402990

**Authors:** Marco Geymonat, Anatole Chessel, James Dodgson, Hannah Punter, Felix Horns, Attila Csikász Nagy, Rafael Edgardo Carazo Salas

**Affiliations:** Pharmacology & Genetics Departments, University of Cambridge, Downing Street, Cambridge, CB2 3EH, United Kingdom.; The Gurdon Institute, University of Cambridge, Tennis Court Road, Cambridge, CB2 1QN, United Kingdom.; Department of Computational Biology, Research and Innovation Centre, Fondazione Edmund Mach, San Michele all’Adige, 38010, Italy.; Pázmány Péter Catholic University, Faculty of Information Technology and Bionics, 1083 Budapest, Hungary.; The Randall Division of Cell and Molecular Biophysics and Institute of Mathematical and Molecular Biomedicine, King’s College London, SE1 1UL, United Kingdom.

## Abstract

A key feature of cells is the capacity to activate new functional polarized domains contemporaneously to pre-existing ones. How cells accomplish this is not clear. Here, we show that in fission yeast inhibition of cell polarity at pre-existing domains of polarized cell growth is required to activate new growth. This inhibition is mediated by the ERM-related polarity factor Tea3, which antagonizes the activation of the Rho-GTPase Cdc42 by its co-factor Scd2. We demonstrate that Tea3 acts in a phosphorylation-dependent manner controlled by the PAK kinase Shk1 and that, like Scd2, Tea3 is direct substrate of Shk1. Importantly, we show that Tea3 and Scd2 compete for their binding to Shk1, indicating that their biochemical competition for Shk1 underpins their antagonistic roles in controlling polarity. Thus, by preventing pre-existing growth domains from becoming overpowering, Tea3 allows cells to redistribute their polarity-activating machinery to prospective sites and control their timing of activation.

## Main text

### Introduction

Countless aspects of cellular function rely on the capacity of cells to activate multiple functionalized domains at their cortex simultaneously. This phenomenon underlies cellular behaviors such as migration, differentiation or chemotaxis, but how cells assemble new functional cell polarity areas in the presence of pre-existing ones is not fully understood. The cylindrically-shaped fission yeast (*Schizosaccharomyces pombe*) recapitulates this conundrum. Newly born *S. pombe* cells grow in a monopolar fashion only from their pre-existing ‘old end’ (OE), which was inherited from their mother at cell division. This old end becomes growth competent after an event called ‘old end take off’ (OETO). The ‘new end’ (NE) formed at the site of division during septation, albeit polarized, is functionally incapable of growing until later in the cell cycle when “new end take off” (NETO (Mitchison and Nurse, 1985)) happens and cells switch on bipolar growth for the remainder of the cycle. The NETO bipolar growth switch involves a number of proteins (Huisman and Brunner, 2011), including: the Kelch-repeat factors Tea1 and Tea3, the Tea1-interactor Tea4 and the actin-interacting protein Bud6, which are polarity landmarks delivered to cell ends in a microtubule-dependent manner; the actin-nucleating Tea4-interacting formin For3, which in conjunction with the active Rho-like GTPase Cdc42 assembles an array of actin cables at the cell ends that directs there exocytosis; and the DYRK kinase Pom1, which contributes to confine active Cdc42 to cell ends by restricting the localization of the Cdc42 GTPase Activating Protein (GAP) Rga4 to the cell sides.

### Results and Discussion

Although the mechanistic contribution of most *S. pombe* polarity factors is well-studied, much less is known about Tea3 - a protein distantly related to the ERM (Ezrin/Radixin/Moesin) protein family (Arellano et al., 2002). *tea3****Δ*** (*tea3*-deleted) cells are NETO defective, which has led to the suggestion that Tea3 is an activator of polarized growth (Arellano et al., 2002; Niccoli et al., 2003), and looking at Tea3-GFP revealed that as published (Arellano et al., 2002) the protein localizes both at the cell ends as well as at the septum, i.e. areas of the cell where growth occurs. However, on close inspection we found that Tea3’s localization pattern anti-correlates conspicuously with that of polarized cell growth, much more than originally noted (Arellano et al., 2002). This was particularly obvious when we co-expressed Tea3-GFP with the RFP-labelled β-glucan synthase Bgs4 (a marker of growing cell domains (Cortes et al., 2005)) and found that, though always present to some extent at cell ends, Tea3 anti-correlates with Bgs4 accumulation at the cortex throughout the entire cell cycle (Figure 1A left). Specifically, we found that Tea3 cortical enrichment: a) increases at the NE and decreases at the OE following OETO; b) drops at the NE following NETO; and c) rises again at both cell ends at septation (Figure 1A middle and right). In stark contrast with its earlier suggested role as an activator of polarity (Arellano et al., 2002), these observations suggested that Tea3 becomes enriched at inactive polarity areas and, therefore, that it could be an inhibitor of polarity.

**Figure 1.**
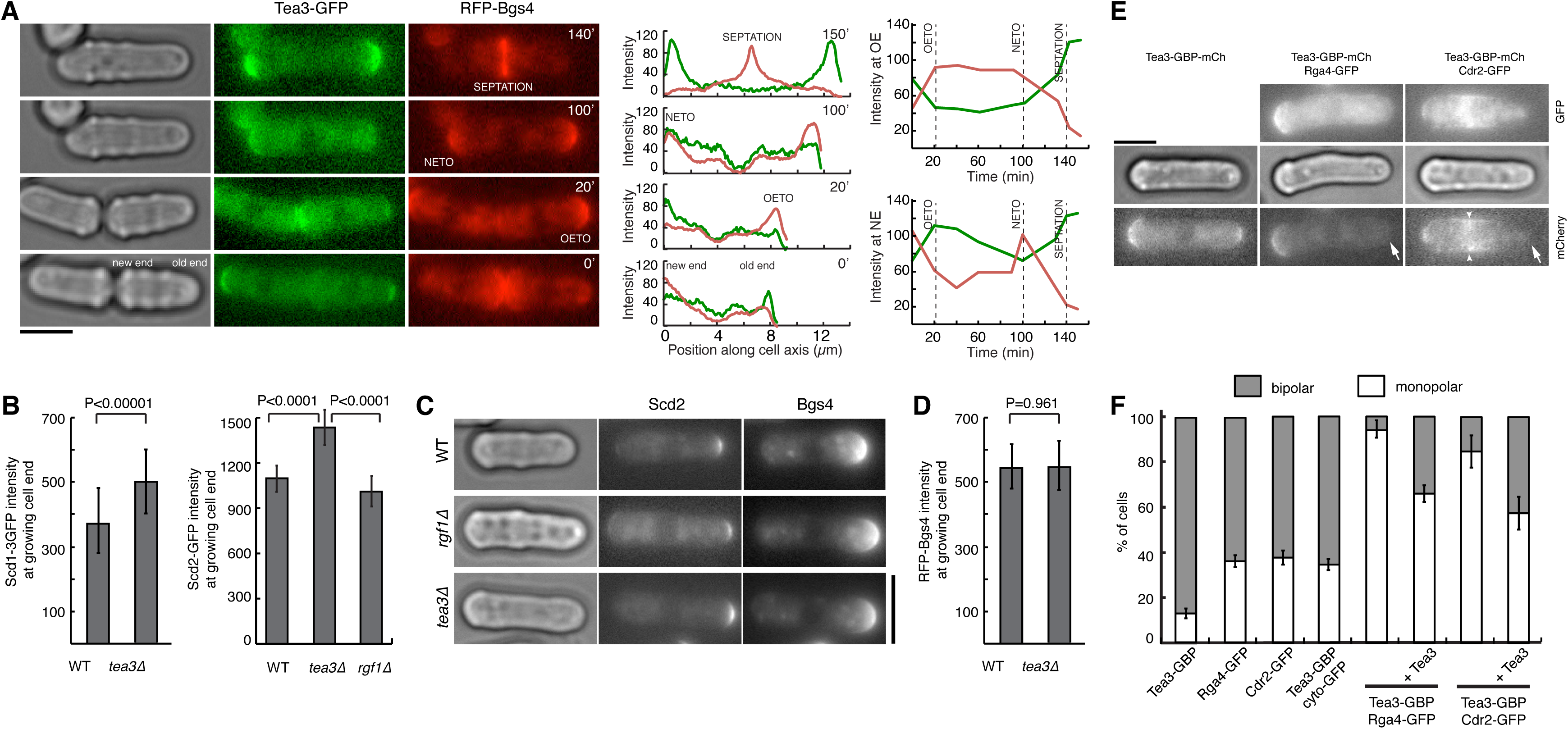
Tea3 is a local inhibitor of Cdc42 activity at growing cell domains. **(A)** Left, A Tea3-GFP RFP-Bgs4 co-expressing cell imaged every 10 minutes and the main polarized growth transitions (OETO, NETO and septation) it undergoes. Middle, Fluorescence intensity profiles of Tea3 (green) and Bgs4 (red) along the cell axis during those transitions. Right, Tea3 (green) and Bgs4 (red) maximal fluorescence intensities at the ‘old end’ and ‘new end’ through the cell cycle. Note that Tea3 counter-mirrors growth. **(B)** Quantification of maximal Scd1-GFP and Scd2-GFP fluorescence at growing end of monopolar wild-type (WT), *rgf1Δ* and *tea3Δ* cells co-expressing Scd1-GFP or Scd2-GFP and RFP-Bgs4cells (n>50 cells/condition). **(C)** Images of monopolar wild-type (WT), *rgf1Δ* and *tea3Δ* cells co-expressing Scd2-GFP and RFP-Bgs4. **(D)** Quantification of maximal RFP-Bgs4 fluorescence at growing end of monopolar wild-type (WT) and *tea3Δ* cells co-expressing Scd1-GFP and RFP-Bgs4cells (n>50 cells/condition). **(E)** Images of cells co-expressing Tea3-GBP-mCherry and Rga4-GFP or Cdr2-GFP. Arrows denote lack of Tea3-GBP-mCherry at the new cell end in the presence of Rga4-GFP or Cdr2-GFP; arrowhead denotes relocalisation of Tea3-GBP-mCherry to the cell middle in the presence of Cdr2-GFP. **(F)** Proportion of monopolar and bipolar cells in exponential cultures with indicated genotypes. Septated cells and cells prior to OETO have not been considered. Average of 2 independent experiments with n>150cells/condition. Error bars represent ± SD. Scalebars: 5μm.

To test this, we looked at the impact of *tea3* deletion (knock-out) on the accumulation of the Cdc42-activating machinery at the cell cortex. The Rho-like GTPase Cdc42 and its GTP-loading co-factors Scd1 (a Cdc42 Guanosine Exchange Factor (GEF)) and Scd2 (a scaffold protein, co-activator of Scd1) are major regulators of polarized growth in this species (Rincon et al., 2014), and the cortical abundances of GTP-Cdc42, Scd1 and Scd2 quantitatively report on the level of polarity activity at cell ends (Abenza et al., 2015; Bendezu et al., 2015; Das et al., 2012). Strikingly, when we measured the levels of Scd1 and Scd2 at the cell cortex in *tea3****Δ*** monopolar cells, we found that both Cdc42 co-factors become significantly enriched at OEs in *tea3****Δ*** cells (Figures 1B-1C and Figure 1-figure supplement 1E-F). This enrichment was specific to *tea3****Δ***, as Scd1/Scd2 did not become enriched at the OEs in monopolar wild-type cells or in the cells of another monopolar mutant *rgf1****Δ*** (Figure 1B, right quantitation; Rgf1 is a Rho1-GEF (Garcia et al., 2006)), and it was specific to Scd1/Scd2, as Bgs4 (itself another positive regulator of polarized growth) did not become enriched at OEs in monopolar *tea3****Δ*** cells (Figure 1D). These data demonstrate that in the absence of Tea3 cell polarity activation at the cortex increases in cells, and indicates therefore that in wild-type cells Tea3’s role is in fact to suppress the enrichment of Cdc42 activators at the cortex and to inhibit Cdc42 activity and cell polarity.

How could Tea3, an inhibitor of polarity, activate polarized growth at NETO?

One possibility would be if, by being enriched at non-growing ends (NEs) before NETO, Tea3 could maintain polarity inhibited there until a signal(s) would lead to its delocalization and, consequently, to the activation of polarized growth at the NE. A prediction of this would be that NETO could only happen once Tea3 becomes displaced from the NE. Therefore we looked in time-lapse sequences whether Tea3 displacement precedes, is coincident with or follows NETO (as assessed by *de novo* RFP-Bgs4 recruitment to the NE; time-lapse interval: 10 min). We found that Tea3 depletion from the NE never precedes NETO (0/11 cells followed by time-lapse) and instead either coincides with NETO (7/11 cells) or follows it (4/11 cells). This suggests that, although we cannot exclude the possibility that Tea3 detachment from the NE could be involved in NETO, if indeed Tea3 inhibits polarity at the NE that inhibition is relieved by NETO rather than causative for NETO. We conclude that the NETO switch likely does not depend on Tea3’s function at the NE.

Could NETO depend on Tea3’s function at the OE (i.e. the pre-existing, actively growing end)? To test this possibility, we sought to deplete Tea3 from the OE. We did this by forcing Tea3 to dimerize with the Cdc42 GTPase Activating Protein Rga4 (normally excluded from actively growing OEs) or with the Wee1 regulating kinase Cdr2 (confined to the cell middle), using the GFP-GBP (<u>G</u>FP <u>B</u>inding <u>P</u>rotein) system (Rothbauer et al., 2008). Co-expression of Rga4-GFP with Tea3-GBP led Tea3 to become depleted from the growing end and both proteins to become enriched at the non-growing end in interphase cells (Figure 1E and Figure 1-figure supplement 1A). This led to a large increase in the percentage of monopolar, NETO-defective cells (Figure 1F and Figure 1-figure supplement 1B). We then co-expressed Cdr2-GFP with Tea3-GBP and found that Tea3 became depleted from both the growing and non-growing ends (Figure 1E and Figure 1-figure supplement 1A), and that this led equally to a large increase in monopolar NETO-defective cells (Figure 1F). Importantly, in both cases additional co-expression of untagged Tea3 partially rescued bipolarity (Figure 1F and Figure 1-figure supplement 1D) without affecting the level of Rga4-GFP or Cdr2-GFP, and hence of Tea3-GBP, at the non-growing end (Figure 1-figure supplement 1C). (Note: the partial rescue is likely due to the fact that Tea3 makes clusters in cells (Dodgson et al., 2013), and has the capacity to self-interact ((Snaith et al., 2005) and Figure 5-figure supplement 1A); therefore it is highly likely that part of the untagged Tea3 interacts with Tea3-GBP and is delocalized from the growth end, and hence cannot fully rescue the NETO defect observed upon co-expression with Rga4-GFP/Cdr2-GFP.) Taken together, these results imply that Tea3 depletion from the growing end impairs NETO, and therefore that the NETO-controlling function of Tea3 is most likely to inhibit Cdc42-mediated cell polarity at the growing OE.

How then could Tea3 control Cdc42 activity? It has been shown that the function of many ERMs and related proteins is controlled by phosphorylation (Fievet et al., 2004; Hirao et al., 1996; Kissil et al., 2002; Nakamura et al., 1995; Pietromonaco et al., 1998; Yonemura et al., 2002) and Tea3 itself is predicted to be a phospho-protein (Beltrao et al., 2009; Carpy et al., 2014; Wilson-Grady et al., 2008). Hence, we reasoned that investigating Tea3’s phospho-regulation *in vivo* could help clarify its function. Immuno-precipitation of Tea3-GFP from wild-type cells revealed one band by Western blot that migrated faster after treatment with lambda phosphatase (Figure 2A lanes 1 and 2), demonstrating that Tea3 is phosphorylated *in vivo*, as predicted. Tea3 phosphorylation was observed in lysates from *cdc10-129* (G1 arrested) and *cdc25-22* (G2 arrested) cells, indicating that it is not cell cycle dependent in any obvious manner (Figure 2A lanes 3 to 6 and Figure 2-figure supplement 1). Interestingly, it was disrupted in cells lacking Mod5, a prenylated protein which concentrates at cell ends and anchors Tea3 cortically at the membrane (Snaith et al., 2005; Snaith and Sawin, 2003) (Figure 2B left panel). Furthermore, we found the phosphorylation to be specifically dependent on the polarity-linked PAK kinases Shk1 and Nak1 (Figure 2C), which also localize at cell ends (Matsuyama et al., 2006).

**Figure 2.**
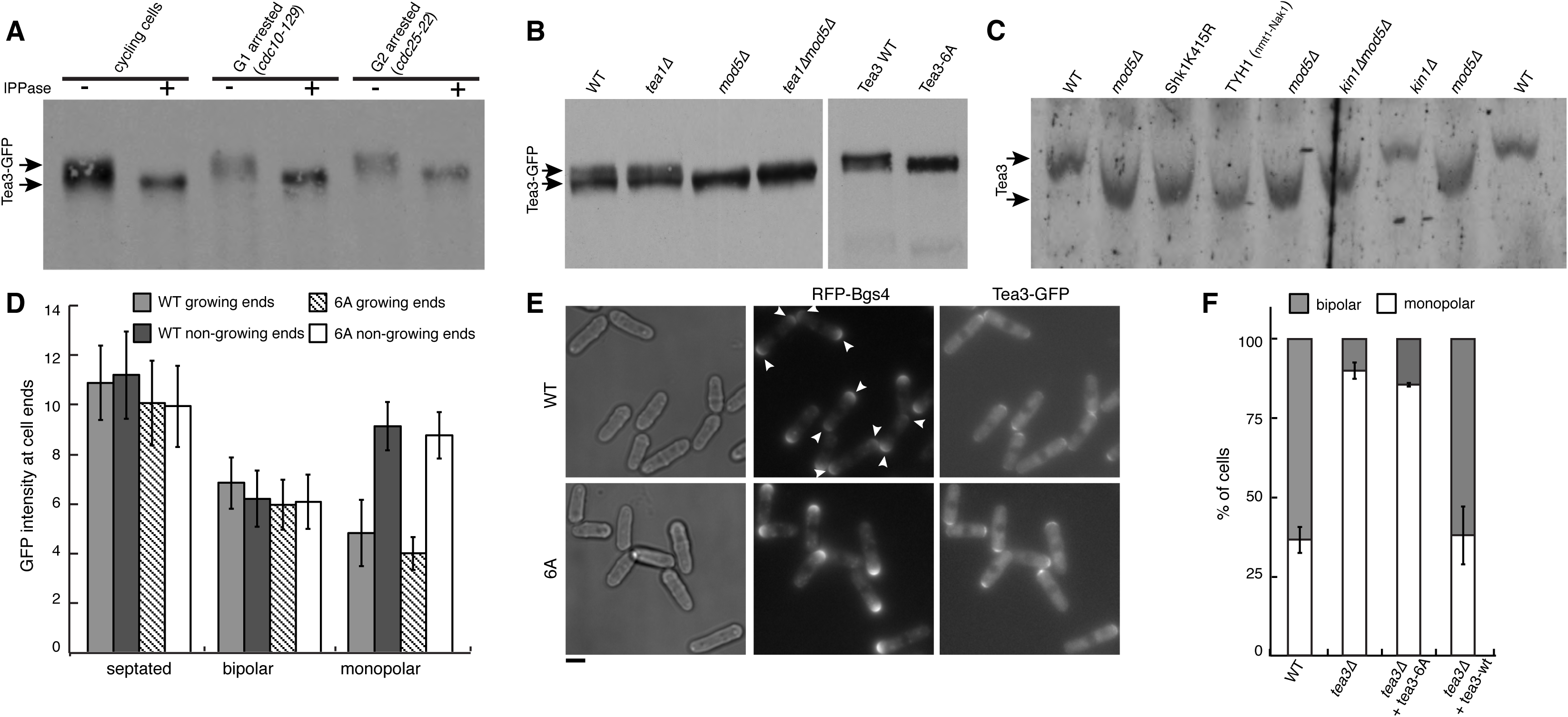
Tea3 function is phosphorylation-dependent and under control by the PAK kinase Shk1. **(A)** Anti-GFP antibody Western blot of Tea3-GFP immunoprecipitated from Tea3-GFP-expressing cycling (left), *cdc10-129* G1-arrested (middle) and *cdc25-22* G2-arrested (right) cells, treated with or without λ-PPase. **(B)** Anti-GFP antibody Western blot of Tea3-GFP from cells of the indicated backgrounds. **(C)** Anti-Tea3 antibody Western blot from cells of the indicated backgrounds. Phos-tag containing 6% acrylamide gel. **(D)** Quantification of Tea3-GFP and Tea3-6A-GFP fluorescence intensity at the cell ends in monopolar, bipolar and septating cells, classified based on their RFP-Bgs4 localization (n>50 cells/condition were measured except bipolar Tea3-6A-GFP cells where n=20, as they were very rare; error bars represent ± SD). **(E)** Example images of cells used for the quantifications in D. Arrows indicate bipolar cells. **(F)** Relative percentages of monopolar and bipolar cells in the cell lines indicated. Average of 2 experiments with n>130 cells/condition. Error bars represent ± SD. Scalebars: 5μm.

These observations imply that Tea3’s phosphorylation is dependent on its cortical anchoring and might take place cortically within cells. Six phosphorylation sites have been found at the Tea3 C-terminus (Beltrao et al., 2009; Carpy et al., 2014; Wilson-Grady et al., 2008) and one of them (position 1045) is a putative PAK consensus site, KRLS (Knaus et al., 1991), similar to the one found in the ERM-related factor Merlin (position 518) and phosphorylated *in vivo* by PAK2 (Kissil et al., 2002). In order to investigate the role of phosphorylation, we generated a phospho-impaired Tea3 mutated in all six sites (Tea3-6A-GFP; Figure 2B right panel). Interestingly, while Tea3-6A-GFP localized normally through the cell cycle (Figure 2D and Figure 2-figure supplement 2) we found that it induces a stark defect in NETO identical to that observed in *tea3****Δ*** cells (Figure 2E-2F). Therefore, Tea3’s PAK-dependent phosphorylation is crucial to its NETO-regulating function.

As the PAK kinase Shk1 has been shown to interact directly with Scd2, and by complexing with Scd2 it has been hypothesized to positively control Cdc42 activity (Bendezu and Martin, 2012; Chang et al., 1999), we surmised that Shk1 might be the relevant kinase at play and that it might interact directly with Tea3 as well. To test that, we expressed and purified Tea3 and Shk1, and tested whether the pure proteins interact. To our good surprise, we found not only that the proteins interact *in vitro* (Figure 3A) but also that Shk1 phosphorylates Tea3 directly (Figure 3B). Given that Tea3 suppresses the cortical enrichment of Scd2 at OEs (Figure 1B), and that both Tea3 and Scd2 are substrates of the kinase Shk1, we wondered if Tea3 might control Scd2 enrichment at the cortex by competitively interacting with Shk1. To ask this, we first expressed and purified Scd2 and we verified that it interacts with Shk1 directly in our experimental conditions (Figure 3C), as reported (Chang et al., 1999). We then tested *in vitro* the binding affinity of Tea3 to Shk1 in absence or presence of excess Scd2, and found that the binding affinity of Tea3 to Shk1 decreases to 30% in the presence of Scd2, demonstrating that Scd2 can outcompete Tea3 (Figure 3D). Reciprocally, we found that the binding affinity of Scd2 to Shk1 decreases to 75% in the presence of excess Tea3, demonstrating that Tea3 can also outcompete Scd2 *in vitro* (Figure 3E). Interestingly, we found that Tea3 outcompetition of Scd2 only occurs in presence of ATP (Figure 3E) and does not happen between excess Tea3-6A and Scd2 (Figure 3F), suggesting a possible differential role for phosphorylation in controlling Tea3-Shk1 versus Scd2-Shk1 interaction.

**Figure 3.**
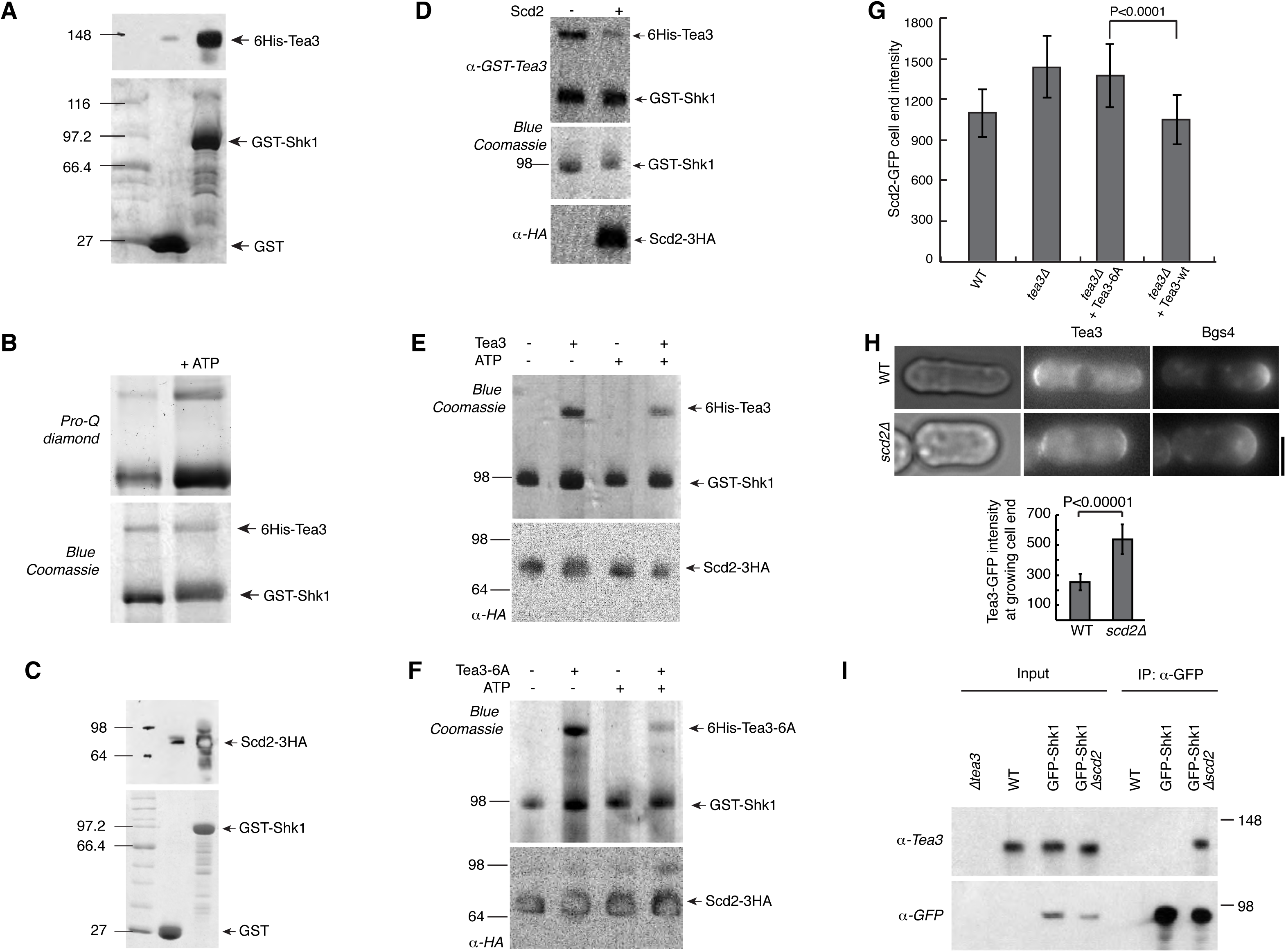
Tea3 is a substrate of Shk1 *in vitro* and it competes with Scd2 for Shk1 binding *in vitro* and *in vivo*. **(A)** Purified 6His-Tea3 binds specifically to GST-Shk1. GST only and GST-Shk1 beads incubated with purified 6His-Tea3 were subjected to SDS-PAGE and Western blot analysis. Upper panel, WB decorated with anti-6His; lower panel SDS-PAGE stained with Blue Coomassie. **(B)** 6His-Tea3 is an in vitro substrate of Shk1. Purified 6His-Tea3 is incubated in absence (left) or presence (right) of 10 mM ATP with purified GST-Shk1. Proteins are subjected to SDS-PAGE and stained before with Pro-Q diamond (Thermo Fisher) to detect phosphor proteins (upper panel) and then with Blue Coomassie (lower panel). **(C)** Purified 6His-Scd2-3HA binds specifically to GST-Shk1. GST only and GST-Shk1 beads incubated with purified 6His-Scd2-3HA were subjected to SDS-PAGE and Western blot analysis. Upper panel, WB decorated with anti-HA; lower panel SDS-PAGE stained with Blue Coomassie. **(D)** Scd2 competes with Tea3 for Shk1 binding. 6His-Tea3/GST-Shk1 complex bound to gluthatione beads is incubated with or without 10x excess of 6His-Scd2-3HA. After extensive washing proteins are analysed by SDS-PAGE and western blotting. Upper panel, anti-Tea3 (note that this antibody recognize also GST), middle panel Blue Coomassie, lower panel anti-HA. **(E)** Tea3 competes with Scd2 for Shk1 binding in an ATP-dependent manner. 6His-Scd2-3HA/GST-Shk1 complex bound to gluthatione beads is incubated with or without 10x excess of 6His-Tea3 in presence or absence of 10 mM ATP. After extensive washing proteins are analysed by SDS-PAGE and western blotting. Upper panel, Blue Coomassie, lower panel anti-HA. **(F)** Tea3-6A cannot compete with Scd2 for Shk1 binding. 6His-Scd2-3HA/GST-Shk1 complex bound to gluthatione beads is incubated with or without 10x excess of 6His-Tea3-6A in presence or absence of 10 mM ATP. After extensive washing proteins are analysed by SDS-PAGE and western blotting. Upper panel, Blue Coomassie, lower panel anti-HA. **(G)** Quantification of maximal Scd2-GFP fluorescence intensity at the growing ends of monopolar cells expressing wild-type Tea3 (Tea3-WT) or Tea3-6A (n=50 cells for each sample). Error bars represent ± SD. **(H)** Tea3-GFP and RFP-Bgs4 localization in WT and *scd2Δ* cells. Quantification of maximal Tea3-GFP fluorescence at growing end of monopolar WT and *scd2Δ* cells (n=35 cells). Error bars represent ± SD. **(I)** Scd2 competes with Tea3 for Shk1 binding in vivo. GFP-Shk1 is immunoprecipitated with nano-trap magnetic beads (Chromo Tech) in WT, GFP-Shk1 and GFP-Shk1 ?scd2 strains. Immunoprecipitated proteins are subjected to SDS-PAGE and Western blotting using anti-Tea3 (upper panel) and anti-GFP (lower panel) to detect Tea3 and FGP-Shk1 respectively.

Taken together, these results demonstrate that Tea3 and Scd2 competitively interact with Shk1 *in vitro*, suggesting they might also recapitulate features of that behaviour *in vivo*. This was confirmed as follows.

A first prediction is that in *tea3-6A* cells, which phenocopy *tea3****Δ***‘s NETO delay (Figure 2E-2F), Scd2 should also become enriched at the OE cell cortex in monopolar cells, given that *in vitro* Tea3-6A cannot outcompete Scd2-Shk1 interaction (Figure 3F). We found that indeed Scd2 becomes significantly enriched at OEs in those cells like in *tea3****Δ*** cells and as observed in *orb2-34* cells (Das et al., 2012) (*orb2-34* is a shk1 mutant allele) (Figure 3G). A second prediction is that, just like Tea3 suppresses Scd2 enrichment at the OE cell cortex (Figure 1B), Scd2 should reciprocally also suppress Tea3 enrichment at the OE cortex. As predicted, we found that indeed in *scd2****Δ*** cells Tea3 becomes cortically enriched (Figure 3H). A third prediction is that it should be possible to observe a differential binding affinity of Tea3 or Scd2 for Shk1 *in vivo*, in presence versus absence of the competitor. In agreement with this prediction, we found that whilst an interaction of Shk1 with Tea3 was undetectable *in vivo* in wild-type cells, in *scd2****Δ*** cells that interaction was readily detected (Figure 3I). These data demonstrate that *in vivo* Tea3 and Scd2 competitively bind their common kinase Shk1.

Tea3’s Shk1-dependent antagonism with Scd2 and its Cdc42 negative regulatory function are reminiscent of negative feedbacks shown to be required for fuelling oscillatory behaviour of Cdc42 activity at the cell cortex, in both budding and fission yeast cells. In *S. cerevisiae*, it was recently demonstrated that a negative feedback provided by the PAK kinase Cla4 inhibits the catalytic activity of the Cdc42 GEF Cdc24 (Kuo et al., 2014). In *S. pombe* the existence of a negative feedback required for NETO was postulated based on mathematical modelling and, interestingly, it was also suggested to involve the similar PAK kinase Shk1 (Das et al., 2012).

We wondered if Tea3 is part of that feedback mechanism and asked whether, like GTP-Cdc42, its level fluctuates/oscillates at cell ends. Hence, we co-expressed CRIB-mCh (an indirect reporter of GTP-Cdc42) and Tea3-GFP in cells, imaged them by time-lapse microscopy and we quantitated the fluctuation of their fluorescence intensity at cell ends by automated image analysis. As previously reported, we observed that about half of wild-type cells display CRIB oscillations between both cell ends (period 535.5±207 seconds, n=202 tracked cells; Figs. 4A top and 4B). Conspicuously, we found that Tea3 also oscillates between the cell ends (period 792±297 seconds, n=202 tracked cells; Figs. 4A bottom and 4C) in approximately half of wild-type cells, suggesting that Tea3 could indeed be linked to the GTP-Cdc42 oscillation gearbox. To test this directly, we quantitated CRIB oscillations in *tea3****Δ*** cells and found that in those cells their period of oscillation is 30% longer than in wild-type cells (Figure 4D). Interestingly, Shk1 suppression has been shown to affect GTP-Cdc42 oscillations similarly to Tea3 (Das et al., 2012). This result demonstrates that Tea3 participates mechanistically in the control of GTP-Cdc42 oscillations in cells, and suggests a key Shk1-mediated role for Tea3 in the negative feedback loop that drives these oscillations. Lastly, we asked whether, despite their difference in period of oscillation, Tea3 and GTP-Cdc42 oscillations are linked. By measuring the cross-correlation between the two oscillations we found that Tea3 and GTP-Cdc42 oscillate within a phase difference of ± π/2 and are therefore linked (Figure 4E). Taken together, our results demonstrate that (although likely not the only component (Das et al., 2015)) Tea3 is integral to the negative feedback mechanism that controls the GTP-Cdc42 oscillations important for the bipolar switch in fission yeast.

**Figure 4.**
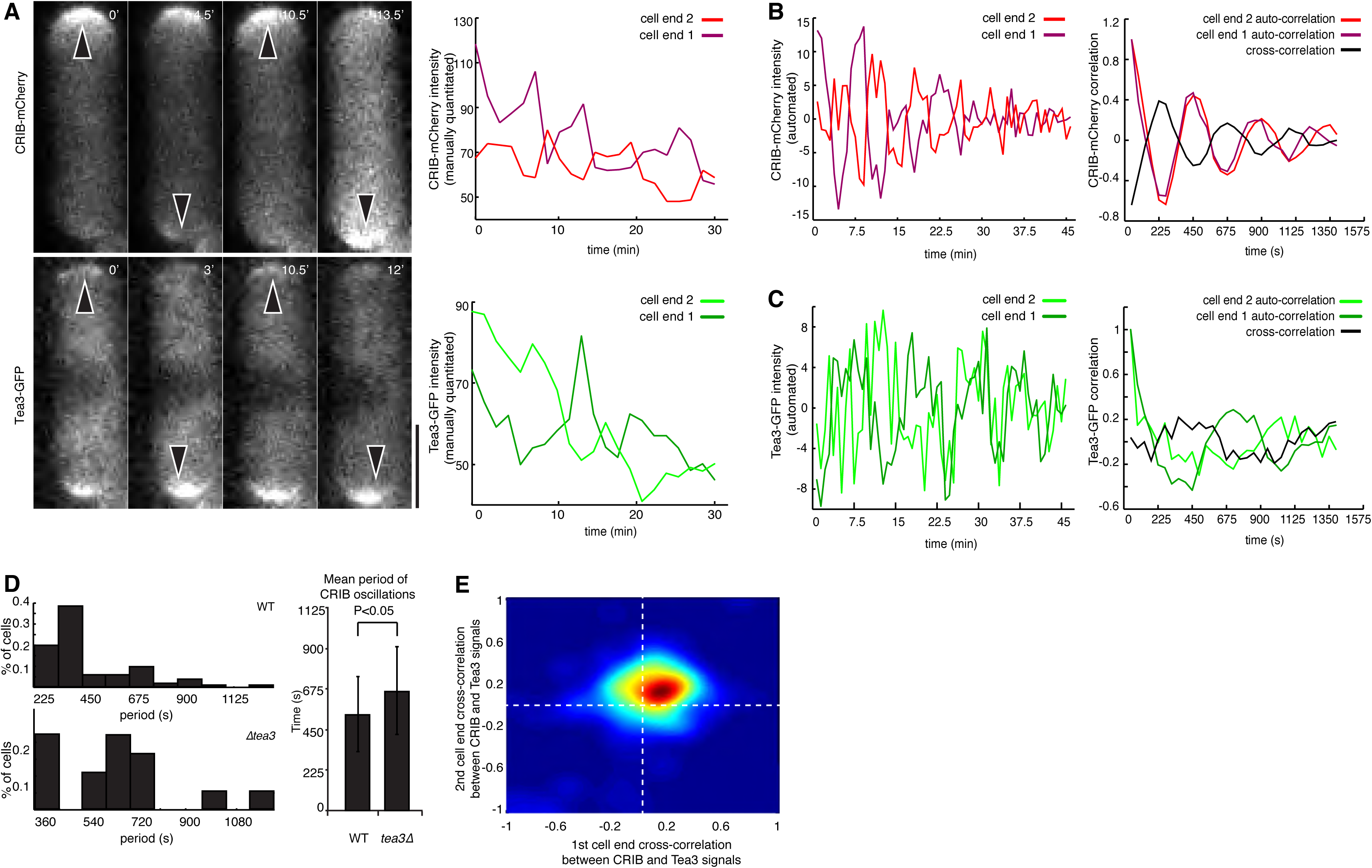
Tea3 is integral part of the mechanism that controls cortical GTP-Cdc42 oscillations in fission yeast. **(A)** Previously described oscillatory behavior of GTP-Cdc42 at cell ends (Das et al., 2012), as observed using a CRIB-mCherry reporter, and newly observed oscillatory behaviour of Tea3, observed via a GFP fusion. Left, timelapse fluorescence images of a CRIB-mCherry Tea3-GFP co-expressing cell (note: 0’ corresponds to the same real timepoint in both top/bottom image sequences). Arrowheads indicate protein enrichment at alternating cell ends. Right, manual quantitation of raw CRIB-mCherry and Tea3-GFP signals from the cell shown on the left. Light and dark coloured lines represent the fluorescence at each of the two cell ends. Note: the CRIB-mCherry and Tea3-GFP image sequences and signals are from the same cell. **(B)** Left, Example of automatically quantitated CRIB-mCherry fluorescence intensity profiles for the two ends of a cell. Right, autocorrelation of each of the two cell end signals (light/dark coloured lines) and cross-correlation between the signals (black line), showing a clear CRIB-mCherry pattern of oscillation between the two cell ends. **(C)** Left, Example of automatically quantitated Tea3-GFP fluorescence intensity profiles from the two ends of a cell. Right: autocorrelation of each of the two cell end signals (light/dark coloured lines) and cross-correlation between the signals (black line), showing a Tea3-GFP pattern of oscillation between the two cell ends. **(D)** Distribution of CRIB oscillation period values in wild-type (top left, n=202) and *tea3Δ* (bottom left, n=56) cells. Right: statistical significance of the difference in CRIB oscillation period between wild-type and *tea3Δ* cells. **(E)** Density plot of the cross-correlation between the Tea3 and CRIB signals measured in the same cell at opposite cell ends, in a population of n=202 tracked cells co-expressing CRIB-mCherry and Tea3-GFP. Note that the density is not centered around zero (which would signify uncorrelation), indicating that the CRIB and Tea3 signal oscillations in a given cell are coupled. **(F)** Schematic model of the Shk1-dependent, Scd2-antagonizing role of Tea3 in controlling activation of polarized cellular growth by locally inhibiting polarity.

It is interesting to note that though Tea3 negatively regulates Cdc42, GTP-Cdc42 and Tea3 oscillate between the two cell ends with distinct periods. Whilst these two findings could seem superficially at odds with each other, it is important to point out that both proteins are imperfect, stochastic oscillators. This is evident from the original publication describing GTP-Cdc42’s oscillatory behaviour in fission yeast (Das et al., 2012), where only half of cells were seen to manifest clear oscillations. Likewise, it can be seen from our results that both oscillators are imperfect and somewhat erratic (Figs. 4A-4C), and similarly we could only detect Tea3’s oscillation in half of cells. Because of this stochastic nature, it follows that an exact relationship between their oscillatory behaviours is probably impossible in practice. Furthermore, we note that the whole system is likely not entirely described by Tea3 and GTP-Cdc42, but may include other stochastically-oscillating players (see for example (Das et al., 2015)), which could cause counter-intuitive differences in the Tea3 and GTP-Cdc42 periods. Despite this, we can affirm that there is a link between both factor’s oscillations, i.e. their phase difference is not random but constrained between ± π/2, which demonstrates that GTP-Cdc42 and Tea3 oscillations are part of the same machinery.

Finally, we wondered if Tea3’s polarity inhibitor function and its functional antagonism with the polarity activator Scd2 might be sufficient to account for Tea3’s role in both regulating GTP-Cdc42 oscillations and controlling NETO. To test this, we did a simplified one-dimensional mathematical model, where we sought to recapitulate the Tea3-Scd2 antagonism by simulating a generic Cdc42 ‘activator’-‘inhibitor’ pair of activities, freely diffusing but cortically localized and retained at cell ends via interaction with microtubule-transported landmark proteins (Figure 5A; see Materials and Methods) (Csikasz-Nagy et al., 2008). The model is based on an earlier one dimensional reaction-diffusion model of cell polarity regulation (Csikasz-Nagy et al., 2008), shown to match several aspects of fission yeast growth patterns and used as the basis for other fission yeast polarity modelling efforts (Das et al., 2012; Thadani et al., 2011). Though based on Scd2 and Tea3, the activator and inhibitor activities we simulated were intentionally kept generic both because of our limited information on the Tea3-Shk1-Scd2 system and to make the least number of assumptions about the morphogenetic properties at play. In the model, we assumed a substrate-limited polarized growth activator (Act) and a substrate-limited inhibitor (Inh), both of which collected cortically at cell ends in an autocatalytic fashion (ActC and InhC, correspondingly). This is not an unrealistic assumption given that we had previously found that polarity regulators in this species localize to cortical clusters likely generated by protein oligomerization (Dodgson et al., 2013) and that at least in the case of Tea3 we found that it has the capacity to self-interact (Figure 5-figure supplement 1A). Also, we assumed that Inh was a faster diffusing form of the inhibitor that interferes with the autocatalytic activation of Act, and conversely that ActC was a slower diffusing form of the activator that interferes with the autocatalytic stabilization of Inh, closing onto a negative feedback loop (Figure 5A, dotted blunt arrows) with a cross inhibition between fast-and slow-diffusing species. The existence of slow (ActC, InhC) and fast (Act, Inh) diffusing forms of the activator and inhibitor were based on our experimental observation that Scd2 and Tea3 diffuse faster in the cytoplasm than at the cell end cortex (Figure 5-figure supplement 1B). This difference in diffusion rates is a prerequisite for Turing-type pattern forming reactions (Goryachev and Pokhilko, 2008; Turing, 1952).

**Figure 5.**
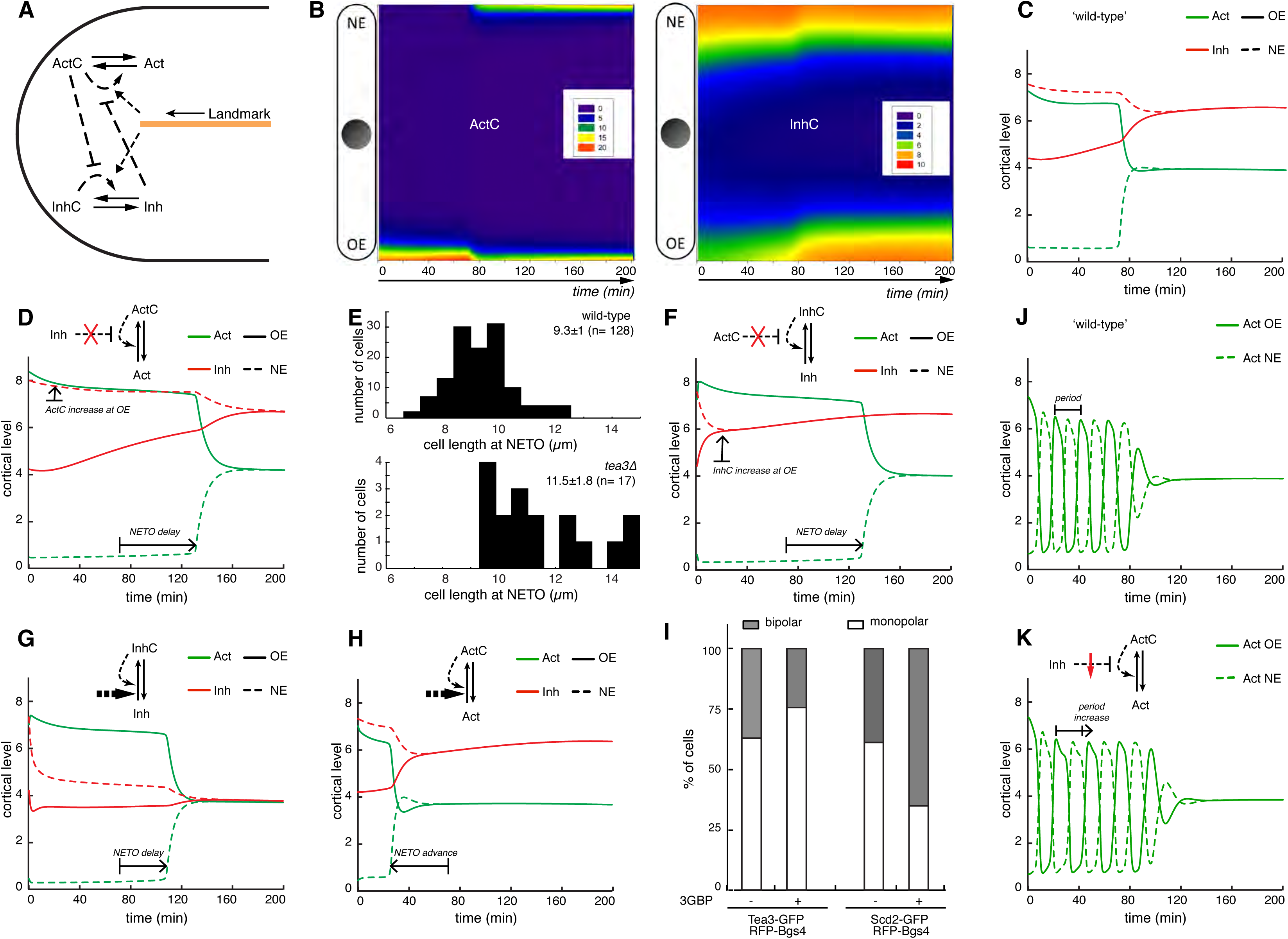
A Cdc42 activator-inhibitor antagonistic system simultaneously accounts for fission yeast polarized growth patterns and Cdc42-oscillations *in silico*. (**A**) An activator–inhibitor model of polarity establishment in fission yeast cells. Cdc42-GTP is controlled by an antagonistic ‘activator’ and ‘inhibitor’ pair, which are freely diffusible and retained at cell ends by microtubule-transported landmarks. Cortical, slow-diffusing activator ActC forms autocatalytically from fast diffusing Act. Similarly, slow-diffusing inhibitor InhC helps its own formation from faster diffusing Inh. The autocatalytic reactions are mediated by polarity ‘landmark’ proteins and ActC inhibits the InhC feedback loop, while Inh inhibits the ActC feedback loop. **(B)** Distribution of ActC and InhC in a 1-dimensional simulated cell. ActC becomes bipolar when the length of the cell reaches 11.3 μm and undergoes NETO. The majority of InhC is localized at the non-growing (i.e. low Act) old end of the cell pre-NETO. OE: old end; NE: new end. **(C)** Average level of total Act and Inh in the 20% outermost region of the old and new ends of the cell in **(B)**. **(D)** Removal of the inhibitory effect of Inh on ActC autocatalysis (*k_4_’*=0min^−1^), as a proxy of *tea3Δ*. Act level increases at the OE and NETO happens when cells reach a longer length of 14.9 μm. **(E)** The experimentally observed size of cells at NETO is statistically bigger in *tea3Δ* cells than in wild-type cells, as predicted by the model (n>200 cells/condition; p-value <0.00005). **(F)** Removal of the inhibitory effect of ActC on InhC autocatalysis (*k_6_’*=0min^−1^), as a proxy of *scd2Δ*. Inh level increases at the OE and NETO happens when cells reach a longer length of 14.7 μm. (**G**) Increase in the background polymerization rate of Inh (*k_5_’*=40min^−1^) causes delay in NETO, happening at a length of 13.9 μm. (**H**) Increase in the background polymerization rate of Act (*k_3_’*=6.9min^−1^) causes advance in NETO, happening at a length of 9.15 μm. (**I**) Mimicked increase in Tea3 and Scd2 polymerization rates by inducing oligomerization of the GFP-labelled proteins, using an oligomer-inducing 3GBP construct confirm the timing of bipolar switch (n=200 cells/condition). (**J**) Perturbation in the degradation rate of Inh (*k_dInh_*=0.15min^−1^) induces oscillations in Act between the two cell ends. (**K**) The period of this oscillation is lengthened if Inh cannot efficiently inhibit Act autocatalysis (*k_4_’*=70min^−1^); the total removal of the effect of Inh would kill oscillations leading to a simulation as on panel (**D**).

We then asked whether this simple activator-inhibitor *in silico* system is sufficient to recapitulate the major features of polarity activation observed *in vivo*.

In a ‘wild-type’ situation, when both the activator and inhibitor were present and allowed to antagonize each other *in silico*, initially small monopolar cells – with the activator concentrated at the OE and consequently the inhibitor concentrated at the NE – grew and became bipolar – with both the activator and inhibitor present in both ends -and this happened at a characteristic ‘NETO’ length (Figs. 5B and 5C), much like what is observed in cells. We first tested what happens when the antagonism between the activator and inhibitor are disrupted *in silico*. When the inhibitor was not able to antagonize the activator, we found that the level of activator at the growing OE of monopolar cells increased (Figure 5D), akin to the enrichment of Scd2/Scd1 observed at the OE in monopolar *tea3****Δ*** cells (Figure 1B). Notably, this enhanced enrichment of activator at the OE coincided with the cells undergoing NETO at a much longer length (i.e. having a NETO delay; Figure 5D) like *tea3****Δ*** cells *in vivo* (Figure 5E), demonstrating that polarity inhibition is required to properly activate new growth areas. Reciprocally, when the activator was not able to antagonize the inhibitor, we found that the inhibitor level at the OE of monopolar cells increased (Figure 5F), similar to Tea3 at the OE in monopolar *scd2****Δ*** cells (Figure 3H), and cells became NETO defective like *scd2****Δ*** cells. (Note: In the model this occurred because in absence of ActC’s effect the inhibitor accumulated at the OE in the clustered form, which is incapable of inhibiting polarized growth there, leading to a NETO defect. *In vivo* this effect could add to the delay caused by the reduced Cdc42 activation in the absence of Scd2 (Kelly and Nurse, 2011), which was not explicitly modelled here.)

We then went on to test various perturbations on the NETO regulating system. We first tested the effect in our *in silico* experiments of perturbations in the cluster forming capabilities of Inh and Act. We found that those showed surprisingly different results in the simulation: when the background polymerization rate of the inhibitor was enhanced, we observed that this led to a predicted NETO delay (Figure 5G), whilst instead NETO was predicted to be advanced when polymerization of the activator was enhanced (Figure 5H). Strikingly, when we mimicked this by enhancing clustering of Tea3 or Scd2 in cells, by inducing oligomerization of the GFP-labelled proteins using a previously published oligomer-inducing 3GBP construct (Dodgson et al., 2013), we found that this led correspondingly to NETO delay or advance *in vivo*, in agreement with the model’s predictions (Figure 5I).

Since our model contains a negative feedback loop (Figure 5A), which could induce oscillations (Novak and Tyson, 2008) we then wondered if we could perturb the *in silico* system to display oscillations in the level of the activator between both cell ends ((Das et al., 2012) and Figure 4A). We found multiple ways to reach this (Figure 5J and Figure 5-figure supplement 2 and see Materials and Methods). Strikingly, when we decreased the effect of the inhibitor on the activator in the simulations (mimicking *tea3****Δ*** cells as above (Figure 5D) we observed longer period oscillations in the activator level at the cell ends, again just like *in vivo* (Figure 4D). We conclude that the antagonism between the polarity activator and the inhibitor suffices to account for all of the basic features of polarity activation observed *in vivo* and explains the mechanistic role of Tea3’s inhibitor role in controlling the activation of new polarized growth at NETO.

In short, in monopolar cells competition of Tea3 and Scd2 for Shk1 binding leads to inhibition of the Cdc42-activating module, allowing the polarity-activating machinery to oscillate between cell ends and enabling a timely NETO switch (Figure 6). We propose therefore that Tea3 is part of the Shk1-dependent negative feedback loop previously found to control NETO (Das et al., 2012). By contrast, in absence of Tea3 GTP-Cdc42 oscillations are impaired and NETO is delayed. Thus, polarity inhibition by Tea3 prevents growing cell ends from becoming overpowering (hyper-enriched) with active Cdc42 and allows its redistribution to prospective growth sites, as required for activating multiple areas of polarity at the cortex contemporaneously (Gierer and Meinhard. H, 1972; Rupes et al., 1999; Turing, 1952) as well as control of the timing of new growth activation. Interestingly, in this model Shk1 acts both at the level of the positive feedback (thought to be mediated by Scd2 (Chang et al., 1999)) and at the level of the negative feedback (mediated by Tea3, shown here), underlying its central role in controlling polarity and the NETO switch.

**Figure 6.**
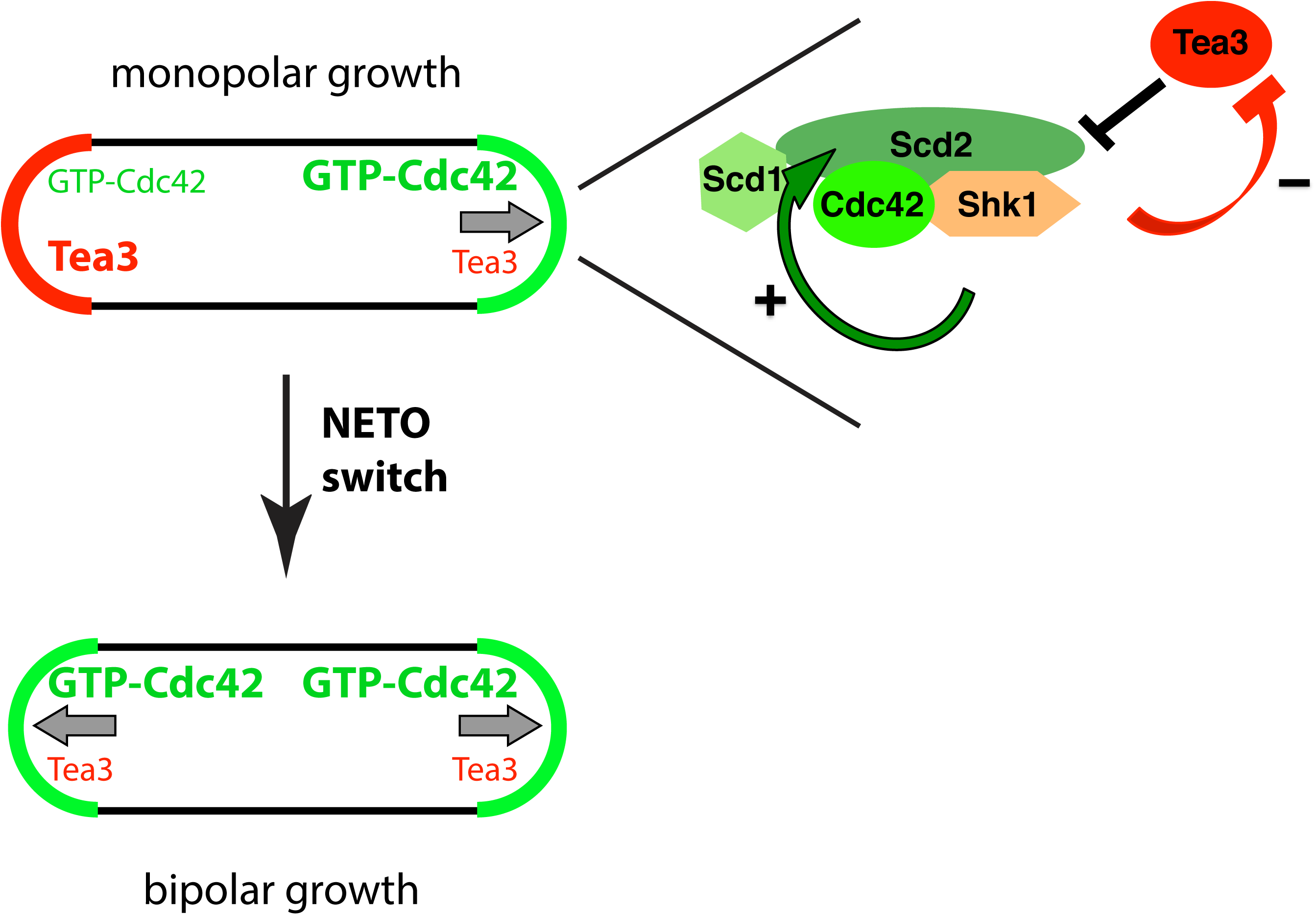
A model of the mechanistic contribution of Tea3 to GTP-Cdc42 oscillations and the bipolar growth switch.

**Table 1.**
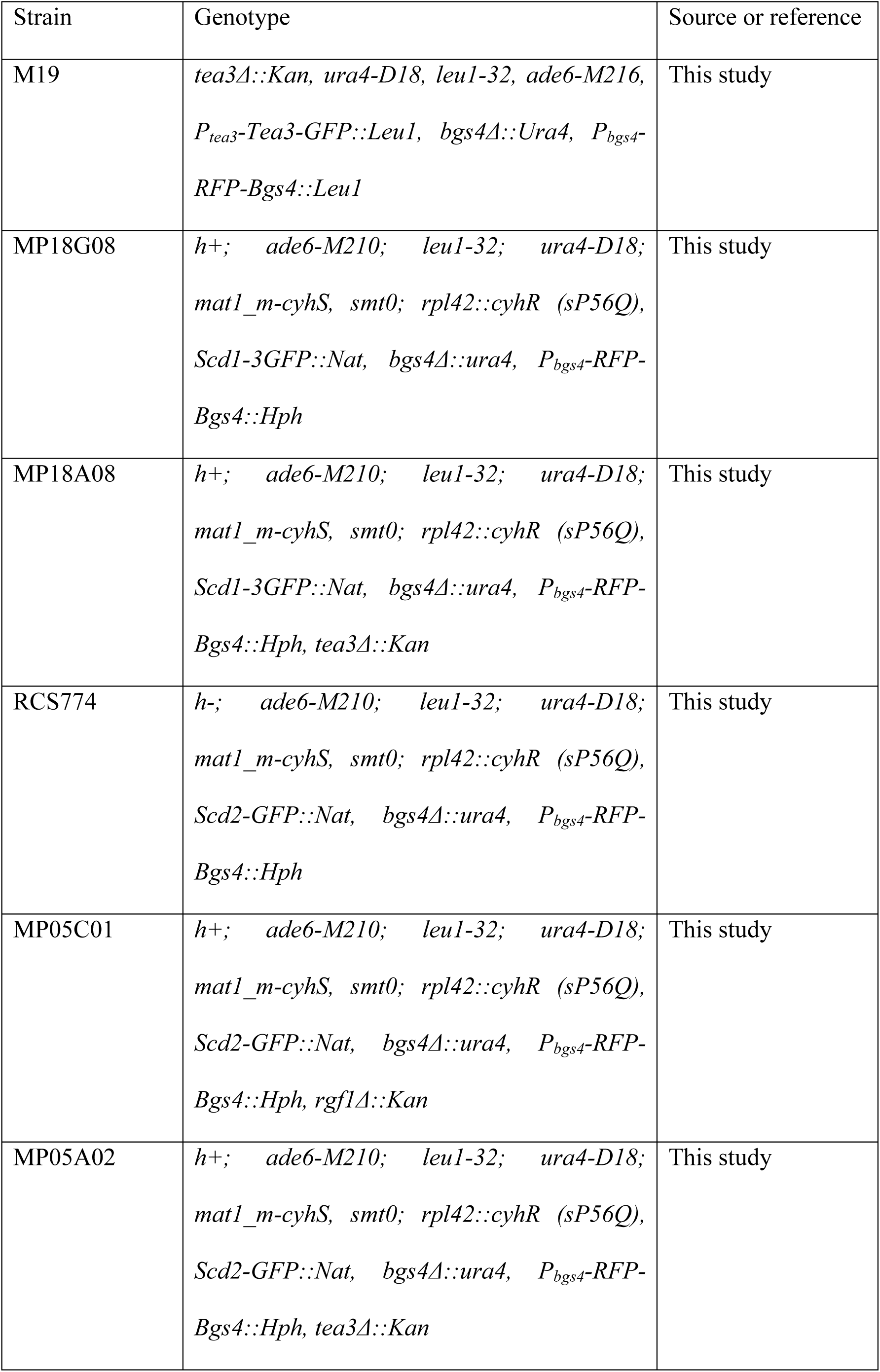
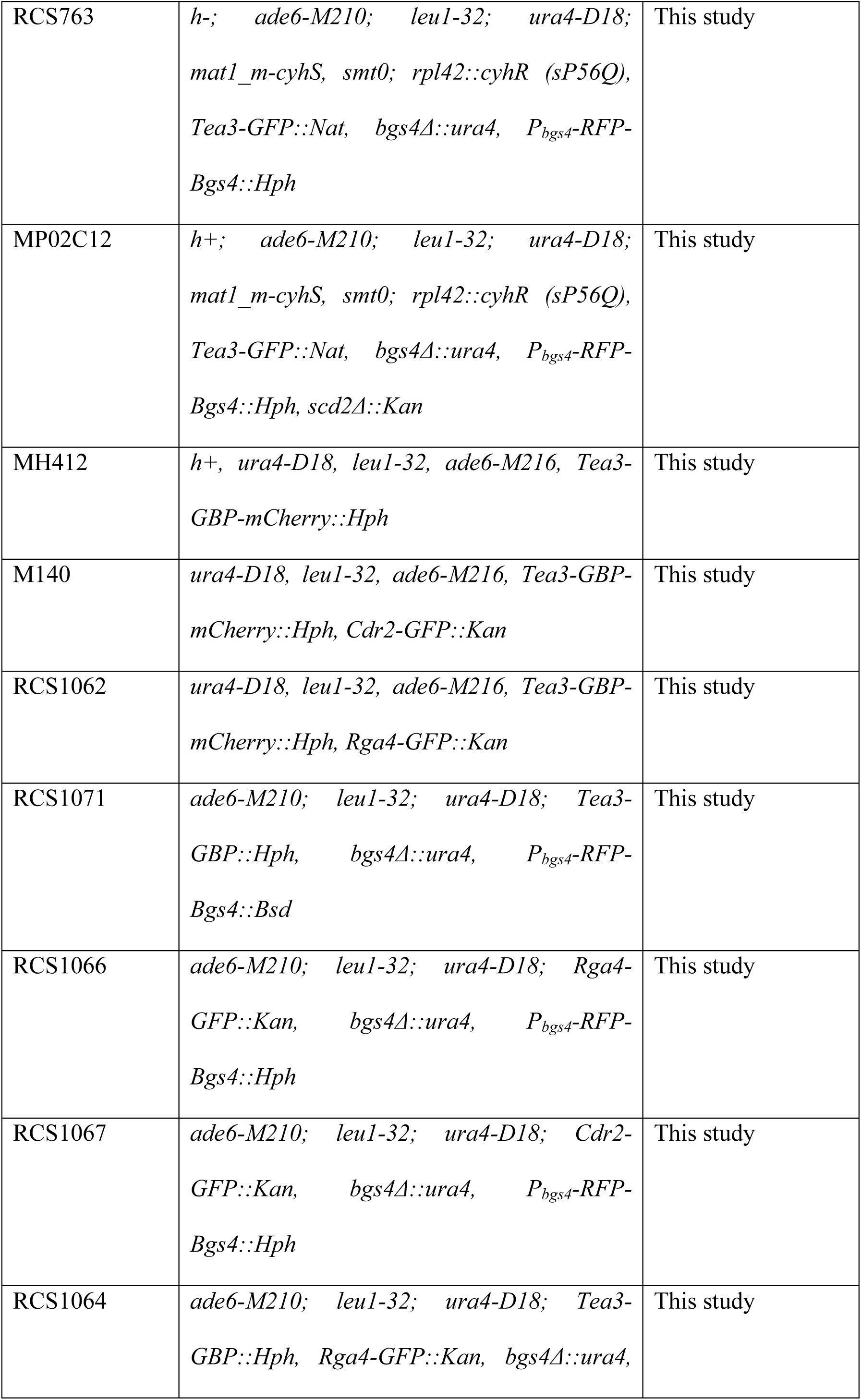
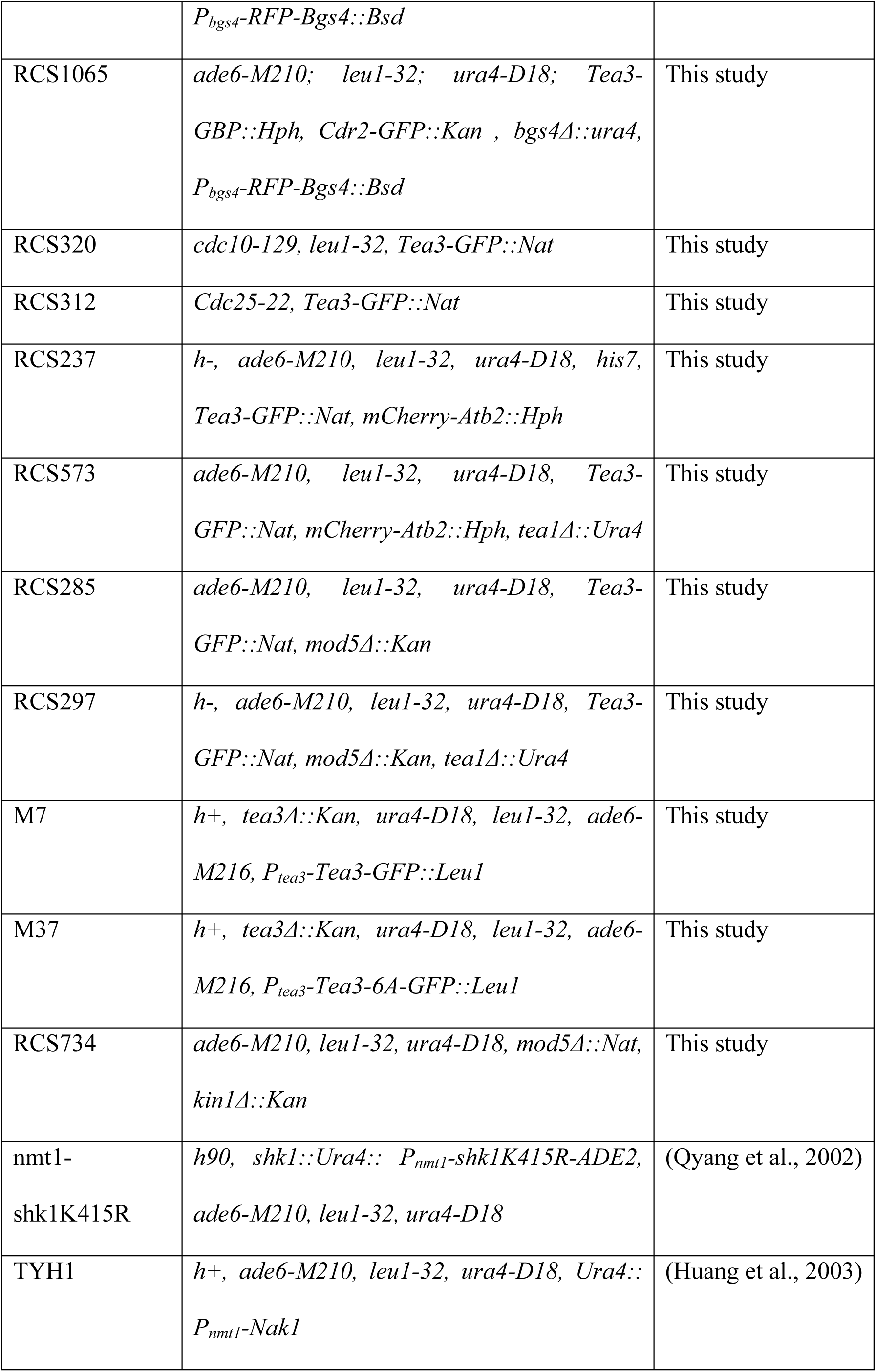
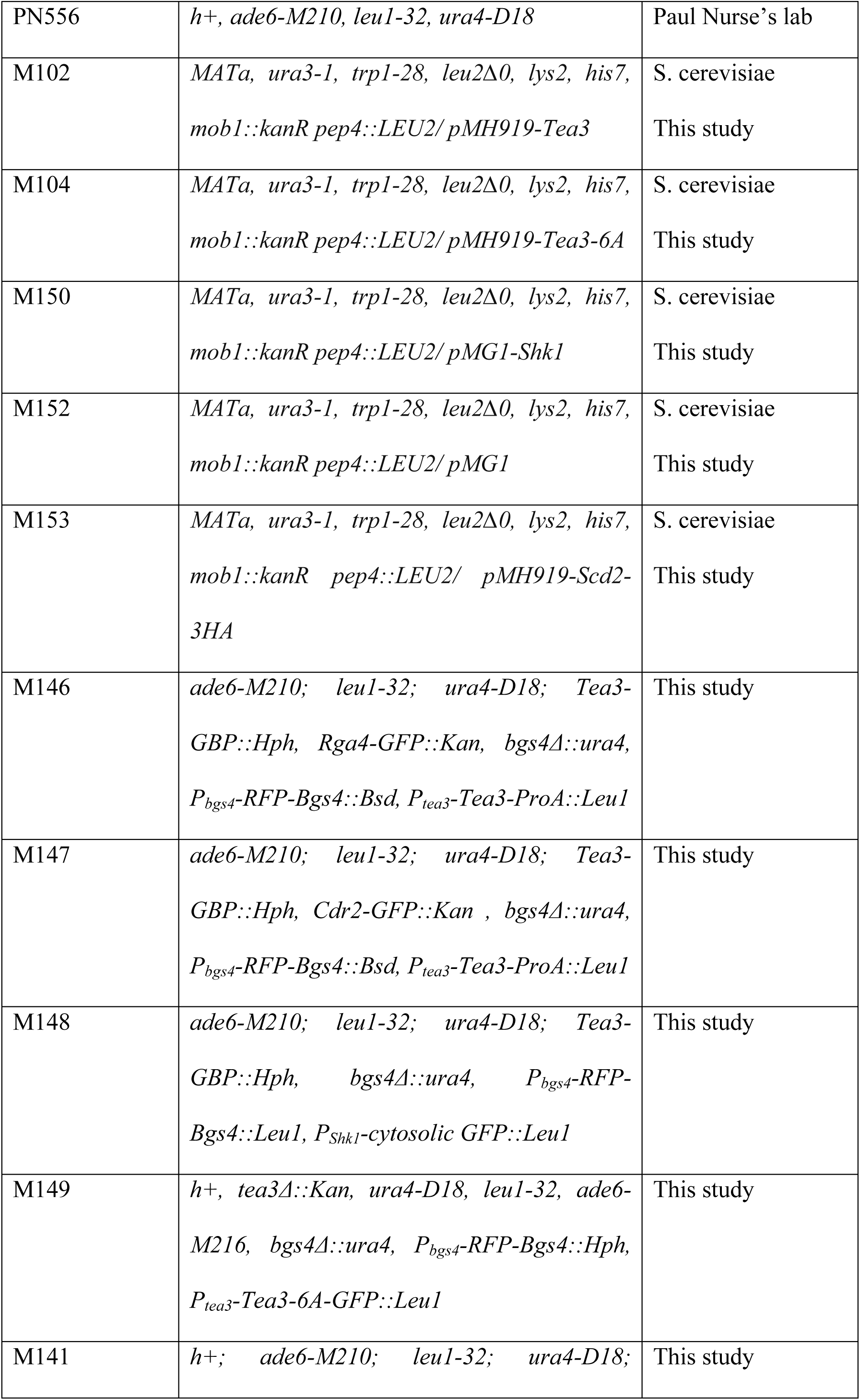
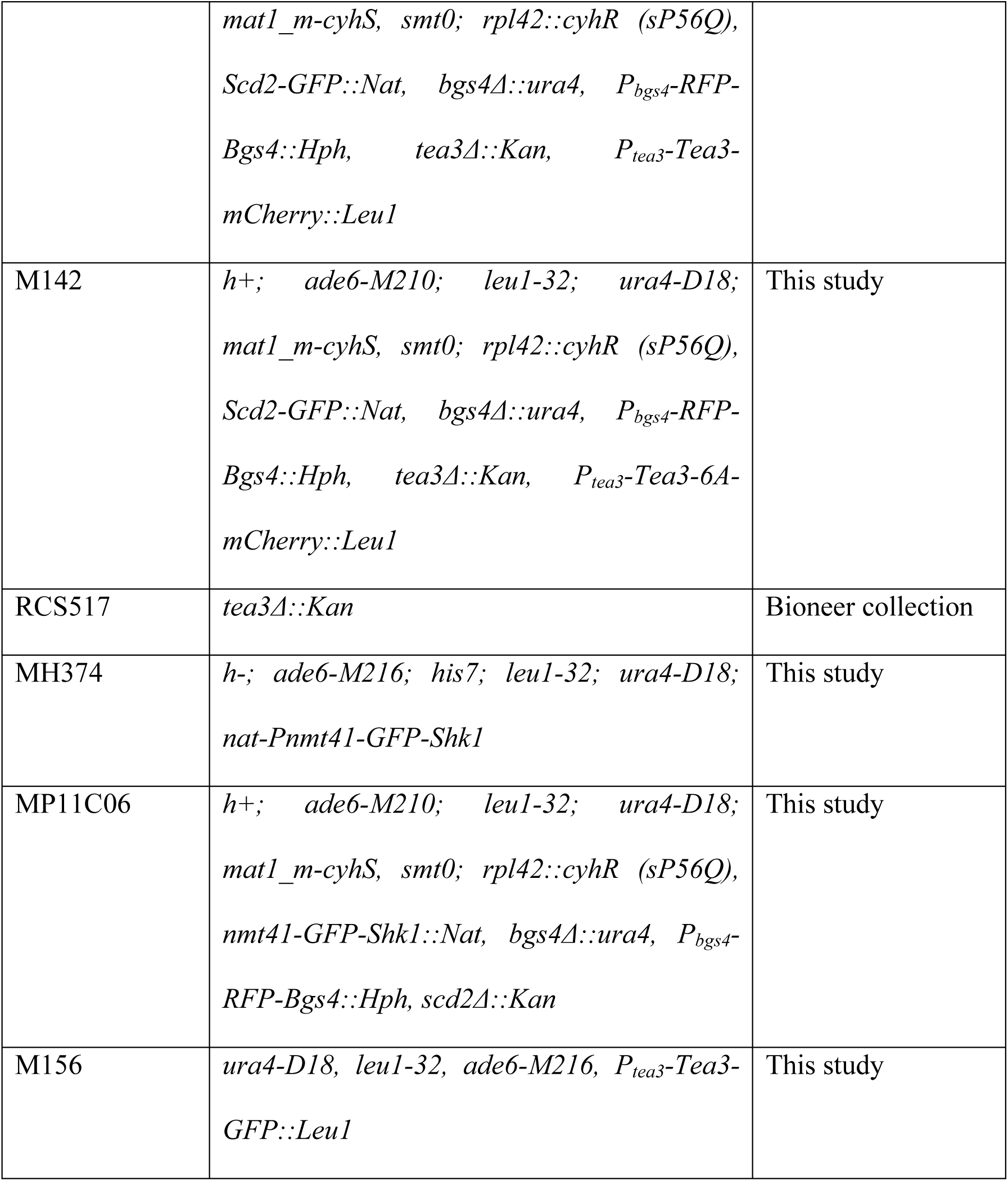

| Strain | Genotype | Source or reference |
| --- | --- | --- |
| M19 | *tea3Δ::Kan, ura4-D18, leu1-32, ade6-M216, P<sub>tea3</sub>-Tea3-GFP::Leu1, bgs4Δ::Ura4, P<sub>bgs4</sub>-RFP-Bgs4::Leu1* | This study |
| MP18G08 | *h+; ade6-M210; leu1-32; ura4-D18; mat1_m-cyhS, smt0; rpl42::cyhR (sP56Q), Scd1-3GFP::Nat, bgs4Δ::ura4, P<sub>bgs4</sub>-RFP-Bgs4::Hph* | This study |
| MP18A08 | *h+; ade6-M210; leu1-32; ura4-D18; mat1_m-cyhS, smt0; rpl42::cyhR (sP56Q), Scd1-3GFP::Nat, bgs4Δ::ura4, P<sub>bgs4</sub>-RFP-Bgs4::Hph, tea3Δ::Kan* | This study |
| RCS774 | *h-; ade6-M210; leu1-32; ura4-D18; mat1_m-cyhS, smt0; rpl42::cyhR (sP56Q), Scd2-GFP::Nat, bgs4Δ::ura4, P<sub>bgs4</sub>-RFP-Bgs4::Hph* | This study |
| MP05C01 | *h+; ade6-M210; leu1-32; ura4-D18; mat1_m-cyhS, smt0; rpl42::cyhR (sP56Q), Scd2-GFP::Nat, bgs4Δ::ura4, P<sub>bgs4</sub>-RFP-Bgs4::Hph, rgf1Δ::Kan* | This study |
| MP05A02 | *h+; ade6-M210; leu1-32; ura4-D18; mat1_m-cyhS, smt0; rpl42::cyhR (sP56Q), Scd2-GFP::Nat, bgs4Δ::ura4, P<sub>bgs4</sub>-RFP-Bgs4::Hph, tea3Δ::Kan* | This study |
| RCS763 | *h-; ade6-M210; leu1-32; ura4-D18; mat1_m-cyhS, smt0; rpl42::cyhR (sP56Q), Tea3-GFP::Nat, bgs4Δ::ura4, P<sub>bgs4</sub>-RFP-Bgs4::Hph* | This study |
| MP02C12 | *h+; ade6-M210; leu1-32; ura4-D18; mat1_m-cyhS, smt0; rpl42::cyhR (sP56Q), Tea3-GFP::Nat, bgs4Δ::ura4, P<sub>bgs4</sub>-RFP-Bgs4::Hph, scd2Δ::Kan* | This study |
| MH412 | *h+, ura4-D18, leu1-32, ade6-M216, Tea3-GBP-mCherry::Hph* | This study |
| M140 | *ura4-D18, leu1-32, ade6-M216, Tea3-GBP-mCherry::Hph, Cdr2-GFP::Kan* | This study |
| RCS1062 | *ura4-D18, leu1-32, ade6-M216, Tea3-GBP-mCherry::Hph, Rga4-GFP::Kan* | This study |
| RCS1071 | *ade6-M210; leu1-32; ura4-D18; Tea3-GBP::Hph, bgs4Δ::ura4, P<sub>bgs4</sub>-RFP-Bgs4::Bsd* | This study |
| RCS1066 | *ade6-M210; leu1-32; ura4-D18; Rga4-GFP::Kan, bgs4Δ::ura4, P<sub>bgs4</sub>-RFP-Bgs4::Hph* | This study |
| RCS1067 | *ade6-M210; leu1-32; ura4-D18; Cdr2-GFP::Kan, bgs4Δ::ura4, P<sub>bgs4</sub>-RFP-Bgs4::Hph* | This study |
| RCS1064 | *ade6-M210; leu1-32; ura4-D18; Tea3-GBP::Hph, Rga4-GFP::Kan, bgs4Δ::ura4,* | This study |
|  | *P<sub>bgs4</sub>-RFP-Bgs4::Bsd* |  |
| RCS1065 | *ade6-M210; leu1-32; ura4-D18; Tea3-GBP::Hph, Cdr2-GFP::Kan , bgs4Δ::ura4, P<sub>bgs4</sub>-RFP-Bgs4::Bsd* | This study |
| RCS320 | *cdc10-129, leu1-32, Tea3-GFP::Nat* | This study |
| RCS312 | *Cdc25-22, Tea3-GFP::Nat* | This study |
| RCS237 | *h-, ade6-M210, leu1-32, ura4-D18, his7, Tea3-GFP::Nat, mCherry-Atb2::Hph* | This study |
| RCS573 | *ade6-M210, leu1-32, ura4-D18, Tea3-GFP::Nat, mCherry-Atb2::Hph, tea1Δ::Ura4* | This study |
| RCS285 | *ade6-M210, leu1-32, ura4-D18, Tea3-GFP::Nat, mod5Δ::Kan* | This study |
| RCS297 | *h-, ade6-M210, leu1-32, ura4-D18, Tea3-GFP::Nat, mod5Δ::Kan, tea1Δ::Ura4* | This study |
| M7 | *h+, tea3Δ::Kan, ura4-D18, leu1-32, ade6-M216, P<sub>tea3</sub>-Tea3-GFP::Leu1* | This study |
| M37 | *h+, tea3Δ::Kan, ura4-D18, leu1-32, ade6-M216, P<sub>tea3</sub>-Tea3-6A-GFP::Leu1* | This study |
| RCS734 | *ade6-M210, leu1-32, ura4-D18, mod5Δ::Nat, kin1Δ::Kan* | This study |
| nmt1-shk1K415R | *h90, shk1::Ura4:: P<sub>nmt1</sub>-shk1K415R-ADE2, ade6-M210, leu1-32, ura4-D18* | (Qyang et al., 2002) |
| TYH1 | *h+, ade6-M210, leu1-32, ura4-D18, Ura4:: P<sub>nmt1</sub>-Nak1* | (Huang et al., 2003) |
| PN556 | *h<sup>+</sup>, ade6-M210, leu1-32, ura4-D18* | Paul Nurse's lab |
| M102 | *MATa, ura3-1, trp1-28, leu2Δ0, lys2, his7, mob1::kanR pep4::LEU2/ pMH919-Tea3* | S. cerevisiae<br>This study |
| M104 | *MATa, ura3-1, trp1-28, leu2Δ0, lys2, his7, mob1::kanR pep4::LEU2/ pMH919-Tea3-6A* | S. cerevisiae<br>This study |
| M150 | *MATa, ura3-1, trp1-28, leu2Δ0, lys2, his7, mob1::kanR pep4::LEU2/ pMG1-Shk1* | S. cerevisiae<br>This study |
| M152 | *MATa, ura3-1, trp1-28, leu2Δ0, lys2, his7, mob1::kanR pep4::LEU2/ pMG1* | S. cerevisiae<br>This study |
| M153 | *MATa, ura3-1, trp1-28, leu2Δ0, lys2, his7, mob1::kanR pep4::LEU2/ pMH919-Scd2-3HA* | S. cerevisiae<br>This study |
| M146 | *ade6-M210; leu1-32; ura4-D18; Tea3-GBP::Hph, Rga4-GFP::Kan, bgs4Δ::ura4, P<sub>bgs4</sub>-RFP-Bgs4::Bsd, P<sub>tea3</sub>-Tea3-ProA::Leu1* | This study |
| M147 | *ade6-M210; leu1-32; ura4-D18; Tea3-GBP::Hph, Cdr2-GFP::Kan, bgs4Δ::ura4, P<sub>bgs4</sub>-RFP-Bgs4::Bsd, P<sub>tea3</sub>-Tea3-ProA::Leu1* | This study |
| M148 | *ade6-M210; leu1-32; ura4-D18; Tea3-GBP::Hph, bgs4Δ::ura4, P<sub>bgs4</sub>-RFP-Bgs4::Leu1, P<sub>Shk1</sub>-cytosolic GFP::Leu1* | This study |
| M149 | *h<sup>+</sup>, tea3Δ::Kan, ura4-D18, leu1-32, ade6-M216, bgs4Δ::ura4, P<sub>bgs4</sub>-RFP-Bgs4::Hph, P<sub>tea3</sub>-Tea3-6A-GFP::Leu1* | This study |
| M141 | *h<sup>+</sup>; ade6-M210; leu1-32; ura4-D18;* | This study |
|  | *mat1_m-cyhS, smt0; rpl42::cyhR (sP56Q), Scd2-GFP::Nat, bgs4Δ::ura4, P<sub>bgs4</sub>-RFP-Bgs4::Hph, tea3Δ::Kan, P<sub>tea3</sub>-Tea3-mCherry::Leu1* |  |
| M142 | *h+; ade6-M210; leu1-32; ura4-D18; mat1_m-cyhS, smt0; rpl42::cyhR (sP56Q), Scd2-GFP::Nat, bgs4Δ::ura4, P<sub>bgs4</sub>-RFP-Bgs4::Hph, tea3Δ::Kan, P<sub>tea3</sub>-Tea3-6A-mCherry::Leu1* | This study |
| RCS517 | *tea3Δ::Kan* | Bioneer collection |
| MH374 | *h-; ade6-M216; his7; leu1-32; ura4-D18; nat-Pnmt41-GFP-Shk1* | This study |
| MP11C06 | *h+; ade6-M210; leu1-32; ura4-D18; mat1_m-cyhS, smt0; rpl42::cyhR (sP56Q), nmt41-GFP-Shk1::Nat, bgs4Δ::ura4, P<sub>bgs4</sub>-RFP-Bgs4::Hph, scd2Δ::Kan* | This study |
| M156 | *ura4-D18, leu1-32, ade6-M216, P<sub>tea3</sub>-Tea3-GFP::Leu1* | This study |

The C-terminal region of Tea3 has a homology with the C-terminal F-actin binding domain present in the ERM-family proteins and Merlin (Arellano et al., 2002; Bretscher et al., 2002). This domain in Merlin and Moesin is subjected to phosphorylation by PAK-like kinases (Hipfner et al., 2004; Kissil et al., 2002) and is essential for their activity. Since Merlin and Moesin have been shown to negatively regulate Rac and Rho respectively (Shaw et al., 2001; Speck et al., 2003) and Tea3 inhibits Cdc42, we propose that there is functional homology between the carboxyl terminal domain of Tea3 and the equivalent domains in the ERM-family proteins. Interestingly, both Merlin and Moesin are at the same time regulators of and regulated by small Rho-like GTPases, and it has been proposed that they are part of a feedback loop important for Rho/Rac regulation (Neisch et al., 2013; Shaw et al., 2001; Speck et al., 2003). We speculate that the role found here for Tea3 of activating polarized growth by inhibiting polarity could extend to other ERM-related proteins. They could equally control spatio-temporally cell polarity plasticity by modulating the tight balance of Rho GTPase activities and turnover at the cortex.

### Materials and Methods

#### S. pombe strains and culture

The *S. pombe* strains used in this study are listed below. Media and general *S. pombe* methods are as described (Moreno et al., 1991).

#### Plasmid construction

A SalI-NotI fragment containing Tea3 ORF and 500 bp of promoter were amplified and cloned into the integrative plasmid pJK148-GFP. This construction, integrated at Leu1 locus, expressed a Tea3-GFP fusion protein that was able to complement the monopolarity of a *tea3Δ* strain. A plasmid expressing a phospho-mutant allele of Tea3 (Tea3-6A) where 6 serine residues (950, 984, 1045, 1058, 1078 and 1080) were mutated in alanine was obtained by several rounds of mutagenic PCR. The mutagenic oligos used were: 5’ CGTAAGCTT**G**CTGAGGTACAAATTGCATTG 3’ (S950A shown in bold, underlined base indicates a silent mutation creating an HindIII site), 5’GCTTCCTCC**G**CTCCCTTGAGATCATACTTT3’ (S984A shown in bold, underlined base indicates a silent mutation eliminating an AflII site), 5’CATAAAAGACTT**G**CTGATGTTATCAACAGTCAGCAAAAATTTTTGTCTTT**G**CCCCACAGGTATCTAAAGAT3’ (S1045A and S1058A are shown in bold, underlined base indicates a silent mutation eliminating an Bsu36I site), 5’CCGCGGGC**G**CATTT**G**C**C**GGCGAAGAAATGCGTGCA 3’ (S1078A and S1080A are shown in bold, underlined base indicates a silent mutation creating an NaeI site). The PCR fragments were cloned into pJK148-Tea3-GFP using BglII-NotI sites. Each mutation was confirmed by plasmid sequencing.

In order to over-express Tea3, the nmt1 promoter (1.1 Kb) was amplified from pREP1 and cloned into a modified version of pJK148 using ApaI-NcoI sites creating pJK148-nmt1. pJK148-nmt1-Tea3NT was obtained by cloning the first 1300 bp (containing a unique EcoRI site) of Tea3 ORF into NcoI-NotI sites of pJK148-nmt1. An ApaI-EcoRI fragment derived from pJK148-nmt1-Tea3NT was then cloned into pJK148-Tea3GFP cut with the same enzymes. Plasmid expressing Tea3-mCherry or Tea3-ProA-tag (i.e. protein A tag) were obtained by substituting GFP with amplified mCherry or ProA-tag using NotI-XmaI restriction enzymes.

#### Lysate preparation and immunoblotting

To assess the phosphorylation status of Tea3 crude extract was obtained using the TCA method described in (Foiani et al., 1994). Briefly, pelleted cells (25 ml at OD_562_< 0.8) were washed with 20% TCA, resuspended in 400μl of 20% TCA and broken with glass beads using a Hybaid Ribolyser (3 cycle of 10 seconds with 3 minutes intervals in ice). 800 μl of TCA 5% was then added and the aqueous extract was spun at 4000 rpm for 10 min. Supernatant was removed and the pellet was resuspended in 100 μl of 1x Laemmli buffer plus 50 μl of Tris 1 M. Tubes were then incubated at 95°C for 5 minutes and spun. Proteins were separated on 6% gels by SDS-PAGE. In order to increase the migration difference between phosphorylated and non-phosphorylated forms of Tea3, 5 μM of Phos-tag (Wako) and 200 μM of MnCl2 were added to the gel as recommended by the manufacturer. To detect Tea3-GFP or untagged Tea3 an anti-GFP antibody (Roche) at 1:1000 dilution or a polyclonal anti-Tea3 antibody (kind gift of P. Nurse) at 1:2000 dilution were used.

#### Immunoprecipitation and phosphatase assay

Cells expressing Tea3-GFP were cultivated overnight at 32 °C in yeast extract with supplements (YES (Moreno et al., 1991); rich medium). Harvested by centrifugation (OD_562_ < 0.8), washed once with cold extraction buffer (EB: Tris 40 mM pH7.5, NaCl 200 mM, KAcetate 50 mM, EDTA 1 mM, MgCl_2_ 2 mM, Triton X-100 0.2%) and resuspended in cold EB containing phosphatase inhibitors (β-glycerophosphate 50 mM, NaF 10 mM and NaO4V 1mM) and protease inhibitors (complete EDTA-free, Roche and PMFS 1 mM). Cells were broken with glass beads using a Hybaid Ribolyser (3 cycle of 10 seconds with 3 minutes intervals in ice). 5 mg of total proteins were mixed with 50 μl of GFP-Trap magnetic beads and incubated at 4 °C for 2 hours. The beads were then washed 6 times with EB and resuspended in PMP buffer containing MnCl_2_ 1 mM (NEB). The beads were split in half and treated or not with 3 μl of λ-PPase (NEB) for 30 min at 30 °C. Beads were washed twice with EB and resuspended in Laemmli Buffer, heated at 95 °C for 5 min and loaded on a gel.

For Shk1-Tea3 co-immunoprecipitation strains expressing GFP-Shk1 were cultivated in YES medium at 32 C and harvested in log phase of growth. Cells were re-suspended in extraction buffer (Tris-HCl 50 mM pH7.5, NaCl 200 mM, Triton X-100 0.1%, glycerol 10%, DTT 2 mM, β-glycerophosphate 50 mM, NaF 10 mM and NaO4V 1mM, protease inhibitors (complete EDTA-free, Roche) and PMSF 1 mM) and broken with glass beads using a Hybaid Ribolyser. 10 mg of total proteins were mixed with 50 μl of GFP-Trap magnetic beads and incubated at 4 °C for 2 hours. The beads were then washed 6 times with washing buffer (Tris-HCl 50 mM, NaCl 150 mM, Triton X-100 0.1% and DTT 5 mM), resuspended in Laemmli Buffer, heated at 95 °C for 5 min and loaded on a gel.

#### Protein purification

6His-Tea3, 6His-Tea3-6A, 6His-Scd2-3HA and GST-Shk1 were expressed in S. cerevisiae (Geymonat et al., 2007). Ni-NTA agarose (Qiagen) and glutathione sepharose 4B (GE Healthcare) were used to purify 6His and GST tagged proteins respectively in accordance with the manufacturer protocols. Purified proteins were dialyzed o/n at 4 C in dialysis buffer (Tris-HCl 20 mM pH 7.5, NaCl 150 mM, DTT 2 mM, glycerol 10%) and stored at −80 C. Typical protein concentration was: for Tea3 and Tea3-6A, 0.5 μg/μl, for Shk1, 0.7 μg/μl and for Scd2, 0.2 μg/μl.

#### In vitro binding and competition

For in vitro Tea3/Shk1 or Scd2/Shk1 binding, strains M152 (expressing GST alone) and M150 (expressing GST-Shk1) were induced for 6 hours. Cells were re-suspended in breakage buffer (Bb: Tris-HCl 50 mM pH 7.5, NaCl 250 mM, Triton X-100 0.1%, DTT 5 mM, glycerol 10 %, EDTA 5 mM and protease inhibitors (complete EDTA-free, Roche and PMFS 1mM)). 10 mg of crude extract from each strain was then incubated with 70 μl of glutathione beads slurry previously washed in Bb without protease inhibitors for 2 hours at 4 C. Beads were then washed 6 times with washing buffer (Wb: Tris-HCl 50 mM pH 7.5, NaCl 250 mM, DTT 5 mM and Triton x-100 0.2%) and one with binding buffer (Bb: Tris-HCl 30 mM pH 7.5, NaCl 150 mM, MgCl2 5 mM and DTT 1 mM). Beads were resuspended in 100 μl of Bb and 5 μg of 6His-Tea3 or 6His-Scd2 were added to each tube. Tubes were incubated in agitation for 1.5 hours at RT then spun and the beads were washed 4 times with washing buffer (Wb2: Tris-Hcl 30 mM pH 7.5, NaCl 150 mM DTT 1mM and Triton X-100 0.1%). Beads were then re-suspended in Laemli buffer and proteins analyzed by SDS-PAGE followed by Western blotting.

For in vitro competition between Scd2 and Tea3 for Shk1 binding, 250 μg of crude extract from strain M150 was used to purify GST-Shk1. GST-Shk1 beads were then incubated in Bb containing 1 μg of Tea3 alone or 1 μg of Tea3 and 10 μg of Scd2. Beads were incubated in agitation for 1.5 hours at RT then spun and washed 4 times with Wb2. Bound proteins were analyzed by SDS-PAGE followed by Western blotting.

For in vitro competition between Tea3 and Scd2 for Shk1 binding, 250 μg of crude extract from strain M150 was used to purify GST-Shk1. GST-Shk1 beads were then washed once in kinase buffer (Kb: Hepes 50 mM pH 7.5, MgCl2 10 mM, MnCl2 1 mM and DTT 1 mM) and then re-suspended in Kb containing 1 μg of Scd2 alone in presence or absence of 10 mM ATP or 1 μg of Scd2 and 10 μg of Tea3 in presence or absence of 10 mM ATP. Tubes were incubated for 45 minutes at 30 C with occasional agitation. Beads were washed 4 times with washing buffer (Wb3: Hepes 30 mM pH7.5, NaCl 150 mM, DTT 1 mM and Triton X-100 0.1%) and proteins were analyzed by SDS-PAGE followed by Western blotting.

#### In vitro kinase assay

In order to detect phosphorylated Tea3 species we used the Pro-Q Diamond phopshoprotein gel stain (Invitrogen). Since the S. cerevisiae purified 6His-Tea3 is slightly phosphorylated and interferes with the Pro-Q staining, Tea3 was pre-treated with λ-PPase (NEB) and then re-purified. For the kinase assay 8 μg of de-phosphorylated 6His-Tea3 and 12 μg of GST-Shk1 were mixed in Kb containing Na3O4V 10 mM in presence or absence of 10 mM ATP. Kinase reaction was carried out for 2 hours at 30 C and then stopped by addition of Laemli buffer and incubation at 99 C for 5 minutes. Samples were analysed on SDS-PAGE and stained with Pro-Q Diamond following manufacturer instructions.

#### Imaging

S. pombe strains were grown at 32 °C to exponential growth and aliquots of 300 μl cells were mounted onto 1.5 coverslip glass-bottom plastic dishes (MatTek; P35G-1.5-14-C) pre-coated with 10 μl 1 mg ml^−1^ lectin. After a 30-min incubation, cells unbound to the lectin-coated glass were removed by washing with medium, and bound cells were kept in a final suspension of 3 ml of medium. Imaging was performed with a DeltaVision System (Applied Precision, USA), based on an Olympus IX81 widefield microscope equipped with a CCD coolSNAP HQ^2^ camera (Photometrix, USA), with a 60x/1.4 N.A. UPLSApo Oil objective. Images were captured and analyzed using SoftWoRx (Applied Precision). Unless otherwise stated, 18 z-stacks with a step of 0.3 μm were filmed with transmitted light and FITC/TRITC filters. Time-lapse images displayed and analyzed in Figure 4 were taken every 45 seconds for 45 minutes.

The monopolarity or bipolarity of exponentially growing cells was determined using RFP-Bgs4 signal at one or both cell ends. Cell in septation or just after cytokinesis (prior to OETO) were not considered. At least 100 cells per condition were analyzed. Error bars represent standard deviation of 2 or 3 independent experiments. T-test was used to compare sets of results.

Quantification of GFP/RFP/mCherry signal was performed using Fiji (http://fiji.sc/Fiji) software after background subtraction. For Tea3-WT and Tea3-6A quantification at the cell ends the sum fluorescence of the cell area of 1.5 μm from the ends have been calculated.

#### Modelling

We extended the original model of Csikász-Nagy et al. (Csikasz-Nagy et al., 2008) with an inhibitor that exists in cortical (*InhC*) and cytoplasmic (*Inh*) forms. Specifically, the equations of the original Csikász-Nagy model were duplicated leading to a system where both *Act* and a newly introduced *Inh* have similar autocatalytic cortical binding reactions that are facilitated by a cortical landmark protein (*u*). The cortical activator (*ActC*, originally named *f* in the Csikász-Nagy model) inhibits the autocatalytic cortical binding of the inhibitor, while the cytoplasmic inhibitor inhibits the autocatalytic cortical binding of the activator (Figure 5A). Thus the only difference between *Act* and *Inh* is in their dynamics and the way they are wired, with the cortical form of *Act* competing with *Inh*’s autocatalysis while the cytoplasmic form of *Inh* competes with *Act*’s autocatalysis. This leads to a situation where a negative feedback loop is introduced in the system ActC --| InhC --| Inh --| ActC. Such systems with three negative effects can induce oscillations (Elowitz and Leibler, 2000) as shown in Figure 5-figure supplement 2.

The model is available as an annotated text file (Geymonat_model_final.ode, Supplementary File) that can be run directly as an.ode file in in the XPPAUT simulation software tool (http://www.math.pitt.edu/∼bard/xpp/xpp.html). In the legends of Figure 5 we provide the parameter changes required to obtain each result reported on Figure 5.

#### Oscillation quantitations and automated microscopy analysis

For the analysis of CRIB and Tea3 oscillations at cell ends, cells were segmented and tracked automatically using in-house algorithms implemented in Matlab. Photobleaching was corrected for each cell for each channel by assuming a constant fluorescence. Mean fluorescence at each cell end for each channel was computed by automatically defining the cell end geometrically, summing the pixels’ greylevel values and dividing by the area. To look at the actual fluorescence fluctuations/oscillations, the continuous trend was removed using empirical mode decomposition (Huang et al., 1998). Assessment of the oscillatory nature of the signal was done in the same way as in (Das et al., 2012): autocorrelation of the signal at each cell end was computed and a given cell end signal was deemed oscillating if, from its starting point of 1, the fluorescence dipped below zero and went back up above 0.2; the period is then the time between two successive maxima. Interestingly, the Cdc42 periodicity of 9 min obtained in this study differs from that describerreported in (Das et al., 2012), where a periodicity of 5 min was described. It is unclear exactly why the oscillation period differs between that study and ours, therefore we can only speculate. One plausible explanation could be the slight differences in the imaging protocols. In particular the temperature used during the imaging experiments might account for this discrepancy, given that all our imaging was done at a Room Temperature of ∼21°C and that paper reports a temperature for all experiments of 25°C. It is possible to imagine biochemical rates of Cdc42 GTPase cycle could be temperature dependent. Other conditions, like optics and cell media, appear comparable and are hence less likely to account for that discrepancy. Importantly, the oscillations of CRIB-mCherry and GFP-CRIB displayed the same period of ∼9min in our experimental conditions (not shown). Hence, it is likely differences in the protocol, and not tagging, that underpins the discrepancy in the period of oscillation between the (Das et al., 2012) study and ours.

For Figure 4E, the cross-correlation plot between the green and red channel was computed for both cell ends for each cell and the density was plotted using kernel density estimation.

To score cell length at NETO (Figure 5E) cells were automatically segmented and tracked, and growth stage was assigned manually, as automated method lacked the required sensitivity given the low number of *tea3Δ* cells undergoing NETO.

#### Sample size & statistical testing

Sample sizes are indicated in all figure legends. p-values were calculated using the t-test function in Microsoft Excel except in Figure 4D, where they were calculated using a two sample Kolmogorov-Smirnov test in Matlab.

### Author Contributions

R.E.C.-S. conceived/led the project and R.E.C.-S. and M.G. designed the general experimental and computational strategy. M.G. carried out all experimental yeast work and imaging, with help from J.D. and H.P. A.C. carried out all quantitative image processing and analysis, with help from F.H. A.C.N. carried out all *in silico* modeling involved. R.E.C.-S. wrote the text with help from M.G. and other co-authors.

## Acknowledgements

We thank E. Piddini, J. Moseley, D. Lew, T. Finegan, J. Pines, K. Sawin, F. Verde, S. Martin, J. Hayles, C. Bradshaw, F. Vaggi and the Carazo Salas group for help and comments, M. Sato, P. Perez, D. Young. P. Nurse and S. Marcus for *S. pombe* strains, P. Zegerman for kind gift of Phos-tag, A. Sossick and N. Lawrence for assistance with imaging, and E. Piddini, D. Lew, J. Mata, J. Moseley and the Carazo Salas group for critical reading of the manuscript. This work was supported by an European Research Council (ERC) Starting Researcher Investigator Grant (R.E.C.-S.; SYSGRO), a Human Frontier Science Program (HFSP) Young Investigator Grant (R.E.C.-S., A.C.-N.; HFSP RGY0066/2009-C), a Biological Sciences Research Council (BBSRC) Responsive Mode grant (R.E.C.-S.; BB/K006320/1) and an Isaac Newton Trust research grant (R.E.C.-S.; 10.44(n)).

---

**Figure 1-figure supplement 1.**
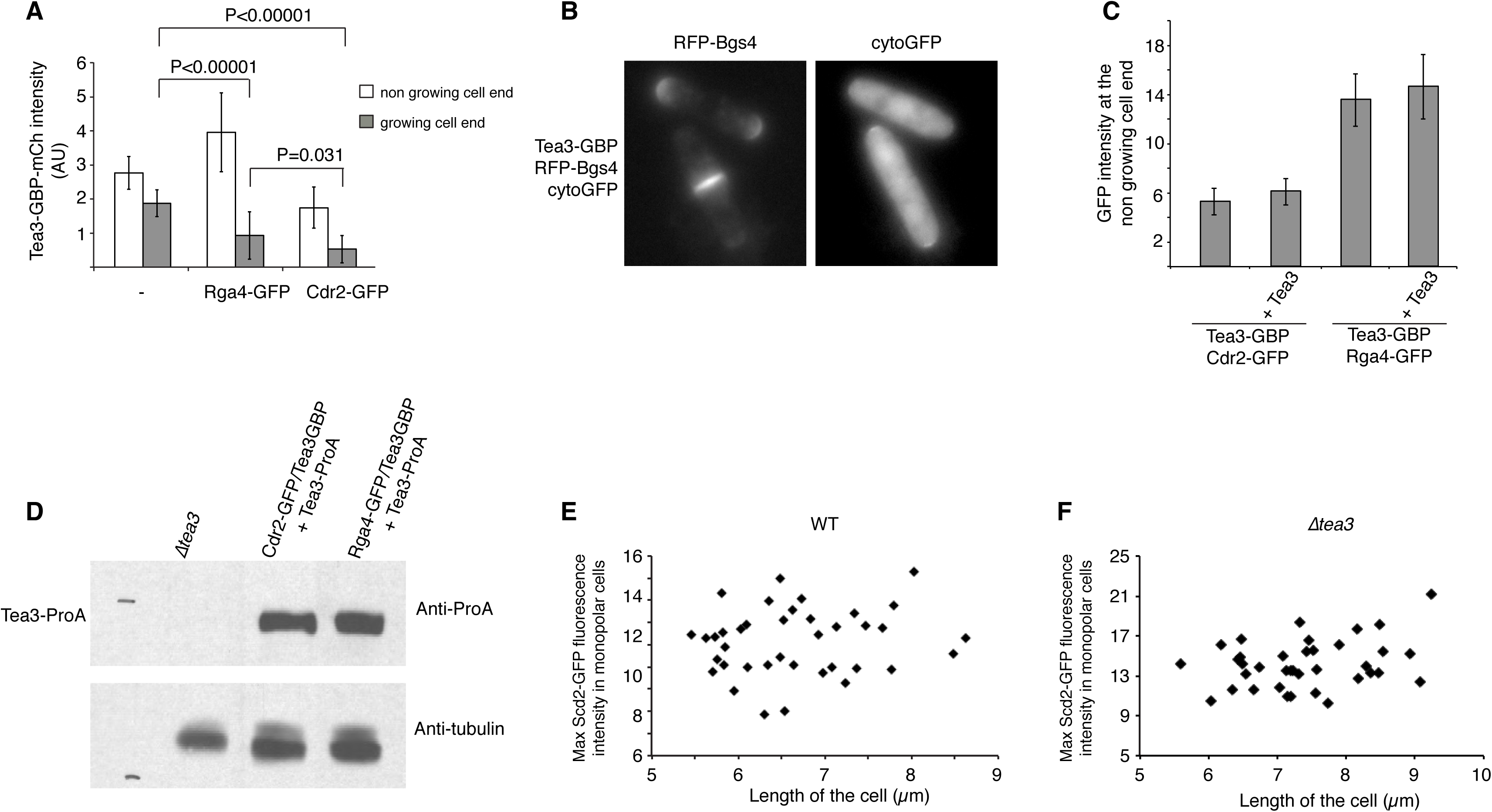
**(A)** Quantification of the Tea3-GBP localisation at the growing and non-growing cell ends in cells expressing Tea3-GBP-mCherry alone or in combination with Rga4-GFP or Cdr2-GFP. n=20 cells per condition. Error bars represent ± SD. **(B)** Images of cells expressing Tea3-GBP and cytosolic GFP. Cell end localisation of the GFP can be observed in cells with a septum, where concentration of Tea3 is high at the cell ends, but not in bipolar cells where concentration of Tea3 at the cell ends is low. **(C)** Quantification of the intensity of Cdr2-GFP and Rga4-GFP at the non-growing cell ends in strains expressing Tea3-GBP alone or in combination with Tea3-ProA. n=20 cells per condition. Error bars represent ± SD. **(D)** Western blot of crude extract derived from strains RCS517, M146 and M147, to show the expression of the Tea3-ProA allele. The upper part of the blot has been probed with a Rabbit Peroxidase anti-Peroxidase antibody (Sigma P1291) and the lower part with a monoclonal anti-tubulin antibody. **(E, F)** Quantification of the intensity of Scd2-GFP in WT **(E)** and *tea3Δ* **(F)** cells in monopolar cells plotted versus the length of the cell. The data demonstrate that there is no increased concentration of Scd2-GFP in longer monopolar cells.

**Figure 2-figure supplement 1.**
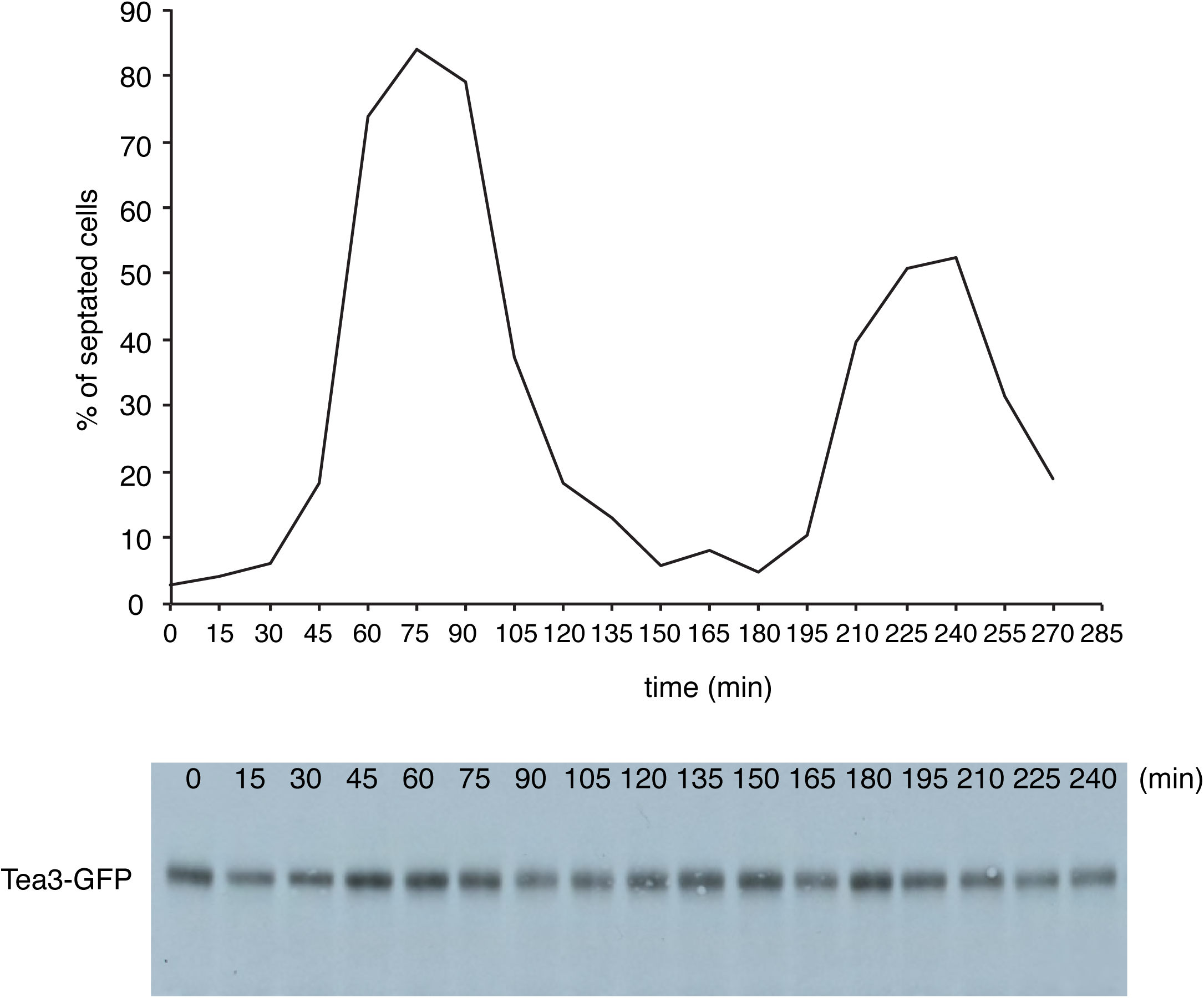
Cell cycle arrest and release of the strain RCS312 (cdc25-22, Tea3-GFP). Cells were arrested for 3h at 37 °C then released at 23 °C. Samples of cells were taken every 15 minutes for TCA extraction and septum staining. Upper panel: septation index during time. Lower panel: SDS-PAGE (6% acrylamide) of crude extract stained with an anti-GFP antibody.

**Figure 2-figure supplement 2.**
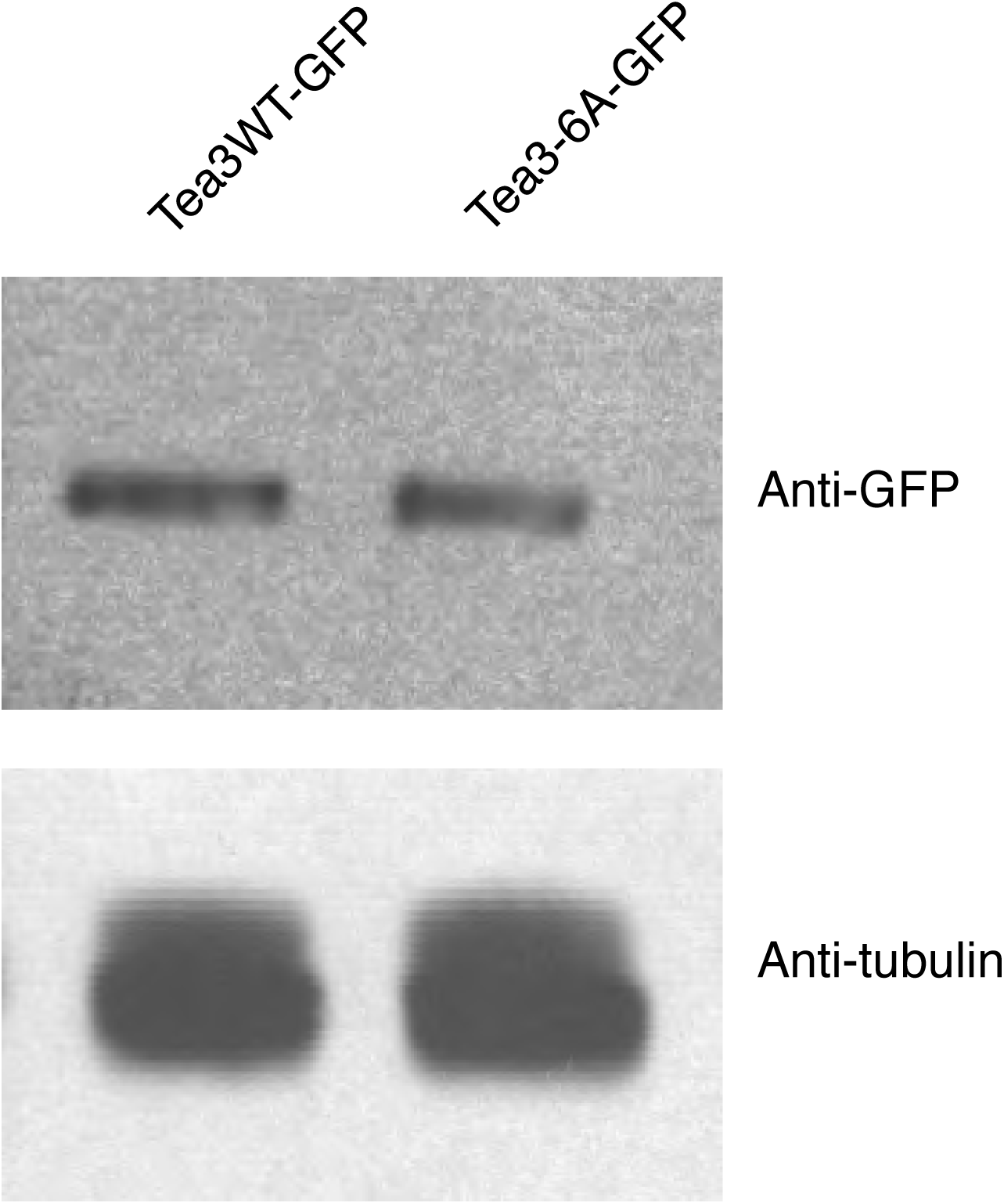
The expression of Tea3WT-GFP and Tea3-6A-GFP is comparable. 70 μg of crude extract from exponentially growing M7 and M37 cells were run on a gel and stained with anti-GFP (upper panel) or anti-tubulin (lower panel) antibody.

**Figure 5-figure supplement 1.**
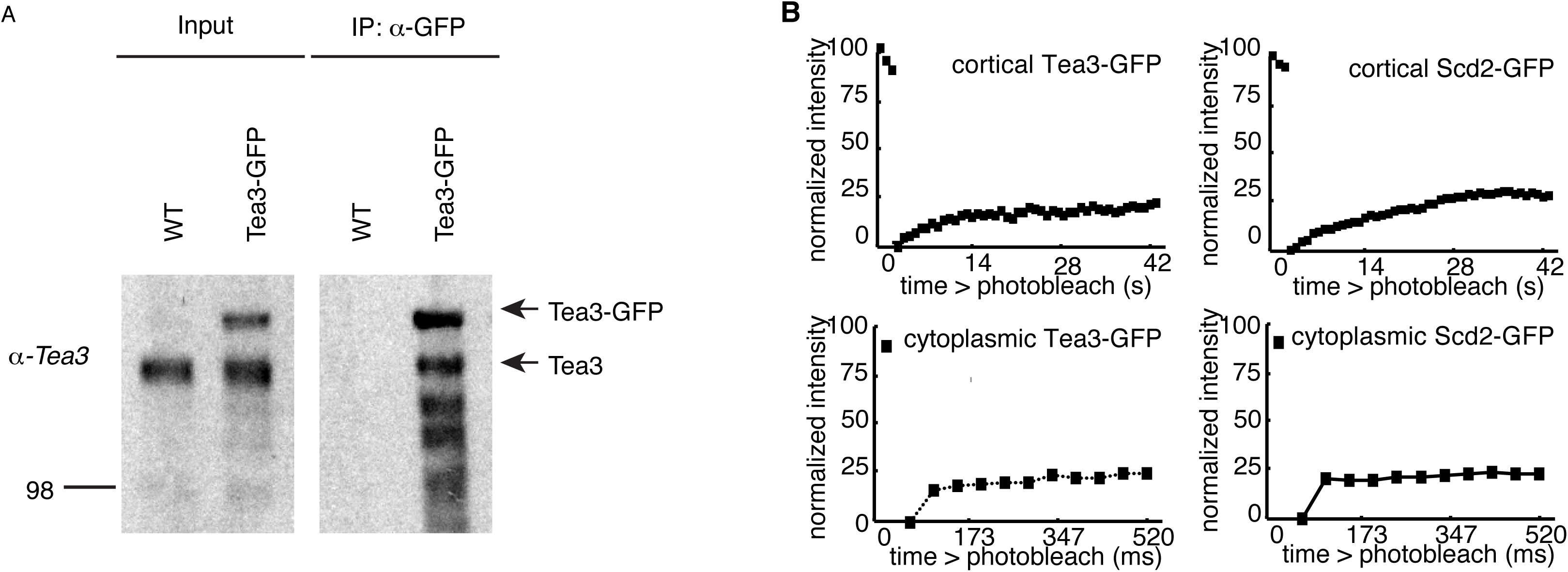
**(A)** Tea3 oligomerises *in vivo*. Tea3-GFP was immunoprecipitated from strains PN556 and M156 with GFP-Trap magnetic beads. After extensive washing bound proteins were analysed by SDS-PAGE and Western blotting using anti Tea3 antibody. Immunoprecipitated Tea3-GFP from strain M156 (expressing also the WT allele of Tea3) is able to co-purify untagged Tea3 demonstrating the ability of Tea3 to oligomerise. **(B)** FRAP experiments using strains RCS763 and RCS774. The photobleached area was half of the cell end (cortical) or inside the cytoplasm (cytoplasmic). For the cortical FRAP, the recovery of fluorescence in the bleached half of the cell end was followed for 1 minute with measurements every 1 second. For the cytoplasmic FRAP, the recovery was followed for 3.7 seconds with measurements every 54 milliseconds. The results show that there are 2 species of Tea3 and Scd2, one highly dynamic in the cytoplasm and another one less dynamic associated to the cortex.

**Figure 5-figure supplement 2.**
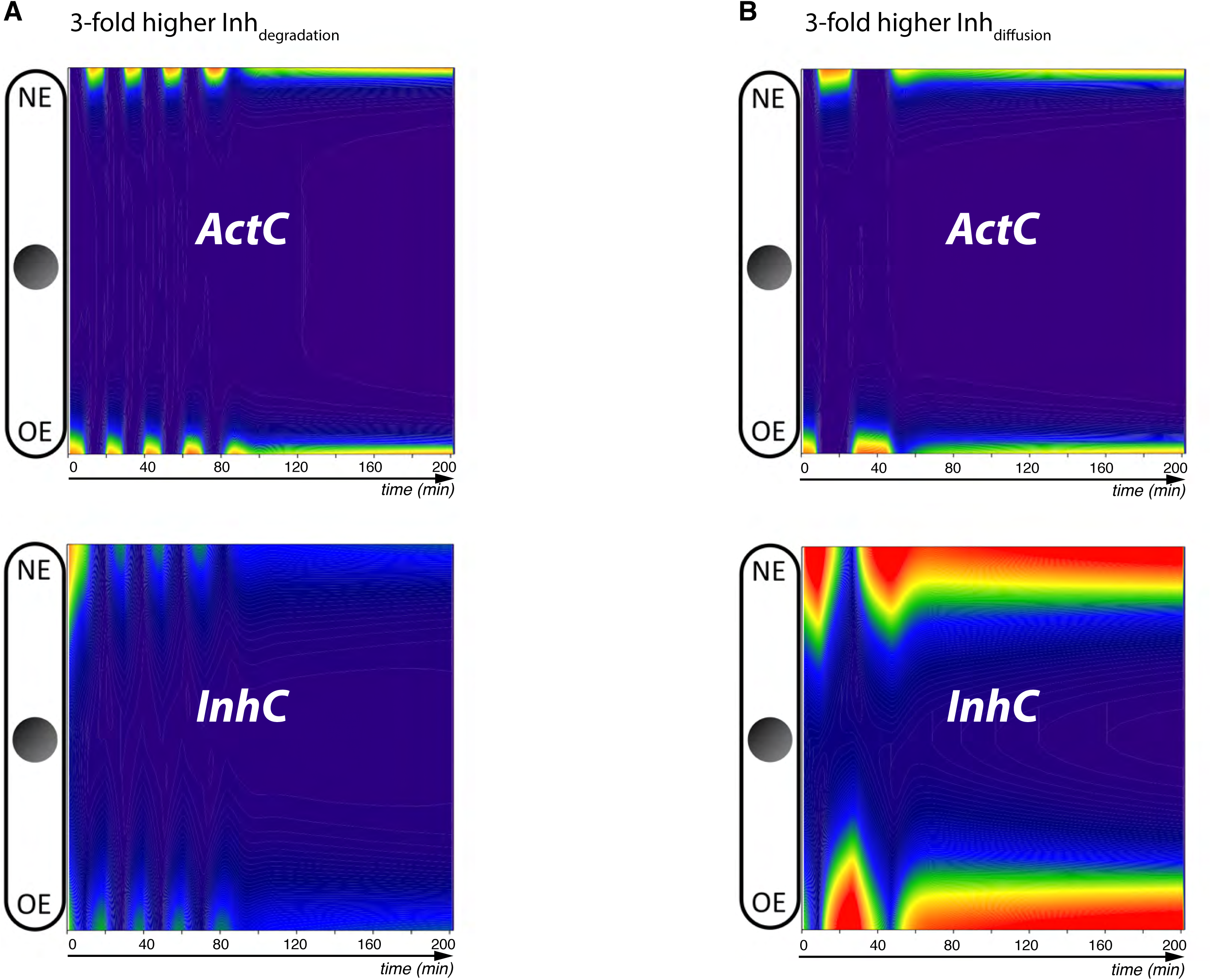
Simulations of oscillations in ActC and InhC levels when the original parameters are perturbed. **(A, B)** Small changes in the degradation (*k_dInh_* = 0.15min^−1^) or diffusion (*D_Inh_* = 240μm^2^/min) rates of the Tea3-like inhibitor ‘Inh’ can induce oscillations in the cortical enrichment of both inhibitor and activator.

